# Browning Lipid Nanoparticles Deliver Metabolic Benefits via Transient Conversion of White Adipose Tissue

**DOI:** 10.64898/2026.09.25.754464

**Authors:** Rohit Sharma, Amanda L Gunawan, Yuchen He, Lin Qi, Nehal Singhal, Jesslyn Lukman, Nayiri Kalindjian, Vyvylyn Tran, Niren Murthy, Andreas Stahl

## Abstract

Therapeutic strategies based on increasing uncoupled respiration (browning) in white adipose tissue (WAT) have great potential for treating obesity-related disorders such as diabetes and steatotic liver disease, but have not had a clinical impact because safe and efficient methods for browning WAT have not been developed. Lipid nanoparticles (LNPs) offer a promising platform for adipose-directed siRNA delivery; however, conventional formulations often exhibit limited activity in adipocytes and can induce inflammatory responses that may restrict repeated administration. Here we demonstrate that LNPs that contain ionizable lipids with three lipid tails and siRNAs targeting TLE3 and ZFP423 can efficiently transfect mature adipocytes and induce browning of WAT *in vitro* and *in vivo* while providing the biocompatibility needed for repeated subcutaneous injections. Repeated local administration of BLNPs in diet-induced obese mice resulted in sustained target gene silencing, increased expression of thermogenic markers, altered energy metabolism, and decreased hepatic lipid accumulation despite minimal effects on body weight relative to scramble siRNA controls. Collectively, these findings establish BLNPs as a promising platform for adipose-directed RNA delivery and targeted modulation of thermogenic pathways, providing a strategy for investigating how localized adipose tissue remodeling influences metabolic physiology.

## INTRODUCTION

Obesity and obesity-related diseases affect billions of people worldwide and are having a devastating effect on global health. For example, 42.5% of adults in the USA are obese^1^ and obesity related diseases such as diabetes, cardiovascular disease and certain cancers are dramatically rising and pose a costly challenge to the American and World healthcare systems^2^. There are currently few treatments for obesity that are safe, effective and have a high rate of patient compliance. Existing obesity treatments such as the GLP-1 agonist semaglutide and bariatric surgery have disadvantages including high cost, injection site effects, loss of lean body mass^3, 4^ and surgery associated complications. Obesity treatments that are safe and effective have the potential to transform healthcare and are urgently needed.

Treating obesity by converting white adipose tissue (WAT) into beige adipose (BeAT) has the potential to generate an effective treatment for obesity with few adverse reactions. Brown and beige adipocytes convert chemical energy from macronutrients to heat^5^ due to the selective expression of uncoupling protein 1 (UCP1), which enables non-shivering thermogenesis. BeAT also modifies metabolism through the release of BAT-derived signals (batokines) such as FGF-21, NRG4, and BMP8b^6^. Consequently, obese patients with high amounts of active brown fat have a healthier metabolic phenotype^7^. Multiple studies in mice and in humans have demonstrated that increased browning and UCP1 expression in WAT are associated with enhanced energy expenditure, improved insulin sensitivity, healthier metabolic profiles, and, in some settings, reduced adiposity and improved glucose homeostasis^8^.

However, despite these promising results, the clinical development of therapeutics that can treat obesity via the browning of WAT has been stagnant because therapeutics that can selectively brown WAT have not been developed^9^. For example, cold exposure, exercise, small molecular activators, and BAT transplantation all have been proposed to stimulate WAT browning and BeAT expansion, but these approaches suffer from issues either with patient compliance, efficacy, and/or safety^10^ and often lack specificity in terms of effects and target tissues^9, 11, 12^. New innovative approaches for WAT browning are needed to define its biological effects and to empower the field to move from animal models into the clinic.

siRNA-based therapeutics have great potential for converting WAT into BeAT in obese patients with minimal side effects because only millimetre volumes of WAT may need to be converted to generate systemic metabolic benefits following localized delivery^13^, which is feasible to achieve after a subcutaneous injection of siRNA. For example, Qiu et al. 2022^14^ demonstrated that siRNA targeting the PRDM16 inhibitor TLE3 complexed to polylysine based lipopolymers, administered subcutaneously once a week for 10 weeks generated metabolic benefits in obese mice. However, although clinically approved siRNA therapeutics have demonstrated that repeated RNA interference can be safe and effective, these approaches have been largely limited to liver-directed applications^15^.

For example, GalNAc-conjugated siRNAs^16^ achieve efficient hepatocyte delivery through the asialoglycoprotein receptor (ASGPR), which is not expressed by adipocytes. Consequently, clinically established strategies for targeted RNA delivery to adipose tissue are lacking, highlighting the need for efficient and biocompatible platforms capable of modulating adipose tissue biology and improving metabolic function. Lipid nanoparticles (LNPs) have great potential for developing therapies that improve metabolic function through adipose tissue remodeling because of their ability to deliver siRNA, low cost, and lower toxicity than polycations and viral vectors. However, relatively little is known about how ionizable lipid structure influences adipocyte delivery, and conventional LNP formulations often induce inflammatory responses that may limit repeated local administration.

Here, we describe browning lipid nanoparticles (BLNPs) containing a three-tailed ionizable lipid (L-36) that support delivery of siRNAs targeting TLE3 and ZFP423 to adipose tissue. Using mouse animal models and human iPSC based microphysiological system (MPS) we show that BLNPs also have the biocompatibility needed for chronic injections of BLNPs. Repeated BLNP administration was associated with sustained thermogenic remodeling of adipose tissue, altered energy metabolism, and reduced hepatic lipid accumulation in diet-induced obese mice.

## RESULTS

### Screening identifies L-36 as an efficient ionizable lipid for adipocyte RNA delivery

To identify lipid nanoparticles capable of efficient RNA delivery to adipocytes, we performed a screening campaign using primary adipocytes to identify ionizable lipid formulations capable of enhancing RNA delivery to adipose cells. The adipocytes used for this screen were derived from the stromal vascular fraction (SVF) of mouse adipose tissue. A commercial library of 30 ionizable lipids, along with the FDA-approved Dlin-MC3 as a benchmark, was screened for their ability to deliver luciferase mRNA to SVF-derived adipocytes (Fig. 1A). Each lipid, designated with an “L,” was formulated into LNPs containing DOPE, cholesterol, and DMG-PEG.

**Fig. 1.**
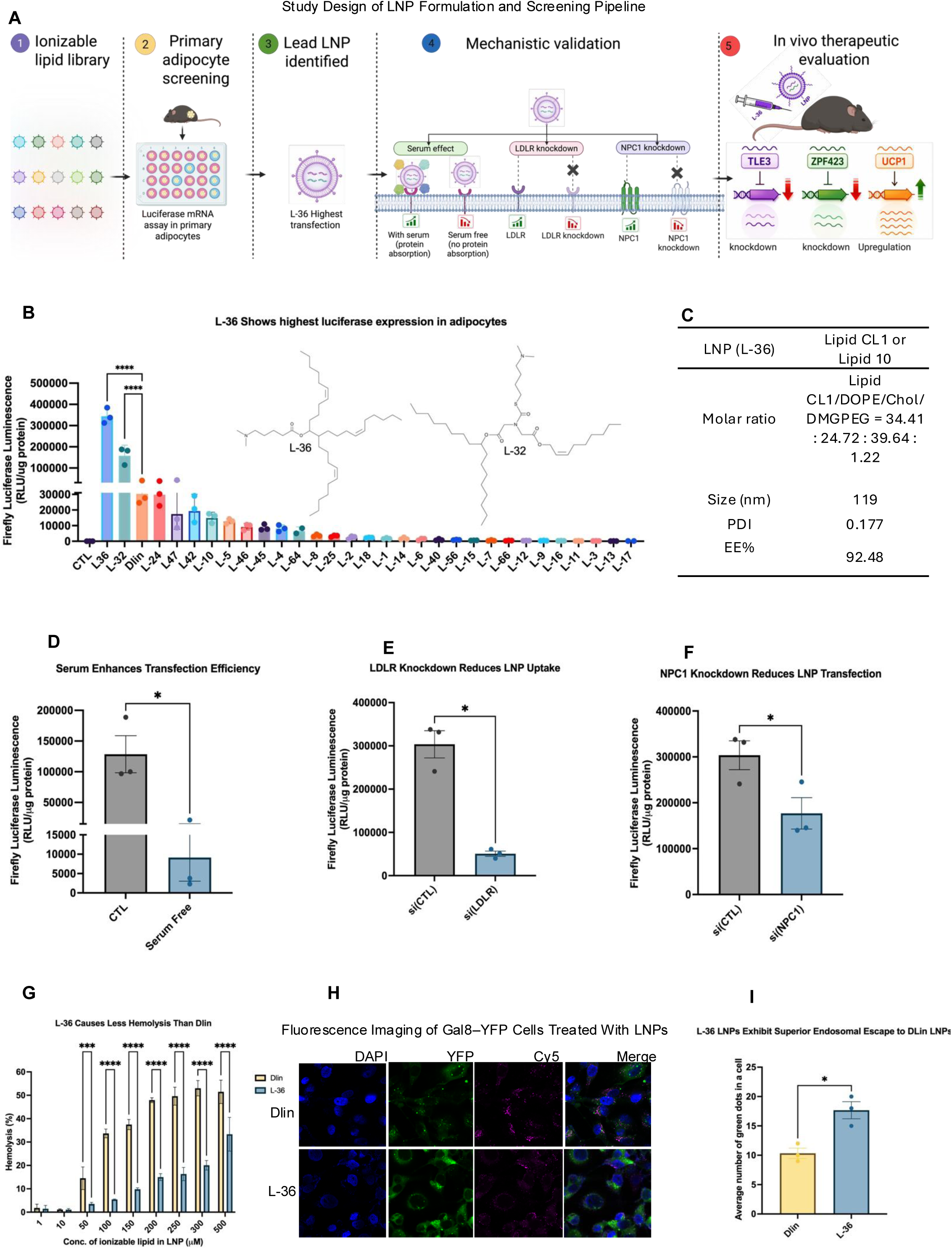
Identification and characterization of L-36 lipid nanoparticles for RNA delivery to adipocytes. **(A)** Schematic overview of the study workflow illustrating the identification, mechanistic validation, and therapeutic application of L-36 LNPs for adipocyte RNA delivery. An ionizable lipid library was screened in primary adipocytes to identify L-36 as the lead LNP for RNA delivery. Mechanistic studies demonstrated that efficient L-36 transfection is promoted by serum, LDLR-mediated cellular uptake, and NPC1-dependent intracellular trafficking. Finally, repeated in vivo administration of L-36 LNPs encapsulating siRNAs targeting TLE3 and ZFP423 induced gene silencing and increased UCP1 expression, establishing the therapeutic strategy evaluated throughout this study. **(B)** Screening of 31 ionizable lipids for luciferase mRNA delivery to murine primary adipocytes identified L-36 and L-32 as the most active formulations. Chemical structures of the two lead candidates are shown. L-36 was selected for subsequent mechanistic and in vivo studies (n = 3). **(C)** Physicochemical characterization of L-36 LNPs, including particle size and polydispersity index (PDI) measured by dynamic light scattering, and RNA encapsulation efficiency determined by the RiboGreen assay. **(D)** RNA delivery by L-36 LNPs under serum-containing and serum-free conditions, demonstrating that serum components facilitate cellular uptake (n = 3). **(E, F)** siRNA-mediated knockdown of the low-density lipoprotein receptor (LDLR; E) and Niemann–Pick type C1 (NPC1; F) significantly reduced luciferase expression, implicating LDLR-mediated uptake and NPC1-dependent endosomal trafficking in L-36-mediated RNA delivery (n = 3). **(G)** Hemolysis assay showing erythrocyte lysis following 1 h incubation with LNPs at 37 °C. L-36 induced substantially less hemolysis than DLin-MC3 across ionizable lipid concentrations of 10–500 μM, indicating improved hemocompatibility (n = 3). **(H)** Representative fluorescence microscopy images of Gal8–YFP-expressing MDA-MB-231 cells treated with DLin-MC3 or L-36 LNPs. The purple channel marks endosomes and lysosomes, whereas the green Galectin-8–YFP signal indicates sites of endosomal disruption. **(I)** Quantification of Gal8 puncta per cell demonstrating significantly greater endosomal escape following treatment with L-36 compared with DLin-MC3 (n = 2 images analyzed per group). Data are presented as mean ± SEM. Statistical significance was determined using an unpaired two-tailed Student’s *t*-test. *P* < 0.05, \*\**P* < 0.001, and \*\*\**P* < 0.0001. Unless otherwise indicated, *n* represents independent biological replicates.

Screening results (Fig. 1B) showed that LNPs formulated with L-32 and L-36 produced significantly higher luciferase expression than the rest of the library and performed better than the FDA-approved MC3-Dlin formulation. Among the tested formulations, L-36 produced the highest luciferase signal and was selected for subsequent characterization. The chemical structure of L-36 (Inset of Fig. 1B) features a distinctive three-tailed hydrophobic architecture, an uncommon architecture among ionizable lipids that may influence nanoparticle behavior and intracellular delivery. L-36 lipid is a next-generation ionizable lipid developed by Genevant, which has shown favorable biocompatibility profile in non-human primates following repeated intravenous administration^17^, supporting its potential utility for repeated nucleic acid delivery applications. L-32 was the second most effective LNP formulation after L-36 in the adipocyte screen. Both L-32 and L-36 contain a single tertiary amine headgroup and unsaturated hydrocarbon tails, but they differ in tail number and saturation pattern (Fig. 1B, inset). L-32 carries two tails, one saturated and one monounsaturated, while L-36 has three monounsaturated tails. This difference likely explains the higher transfection efficiency of L-36, as increasing tail number and distributing moderate unsaturation across all tails can enhance membrane fluidity and endosomal escape during RNA delivery.

Following identification of L-36 as the lead formulation, its physicochemical properties were characterized. L-36 LNPs exhibited a uniform nanoscale particle size with a low polydispersity index (PDI), indicating a homogeneous particle population (Fig. 1C). In addition, the formulation achieved high RNA encapsulation efficiency, demonstrating efficient incorporation of nucleic acid cargo during LNP formulation. These physicochemical characteristics are consistent with a stable LNP formulation suitable for subsequent in vitro and in vivo evaluation.

To investigate structural features associated with adipocyte transfection, we performed a structure-activity relationship (SAR) analysis of the ionizable lipid library based on headgroup composition, degree of unsaturation, number of hydrophobic tails, and tail architecture (Supplementary Fig. S1A– D). Several trends emerged, including improved delivery by lipids containing multiple hydrophobic tails and increased unsaturation, with branched or mixed-tail architectures generally outperforming linear analogues. These analyses support the selection of L-36, whose three-tailed architecture was associated with the highest mRNA delivery efficiency observed in the screen.

To understand how L-36 LNPs achieve efficient mRNA delivery to adipocytes, we examined the L-36 LNP transfection efficiency in serum free conditions, after inhibiting LDLR expression, and after inhibiting NPC1 expression, which is a protein involved in endosomal trafficking (Fig. 1D–F). Luciferase expression decreased significantly under serum-free conditions (Fig. 1D), indicating that serum components facilitate internalization, most likely by forming a protein corona that may enhance receptor engagement. To identify specific receptors involved in uptake, we silenced the low-density lipoprotein receptor (LDLR) and CD36 in differentiated adipocytes using siRNAs. Knockdown of LDLR resulted in a strong reduction in luciferase expression following L-36 LNP treatment (Fig. 1E), which demonstrates that LDLR is the primary receptor mediating internalization. In contrast, silencing CD36 had no detectable effect (Supplementary Fig. S1E), indicating that this scavenger receptor does not contribute to LNP uptake. Importantly, receptor knockdown did not alter the expression of adipocyte differentiation markers, confirming that the observed changes in luciferase expression were due to altered uptake rather than changes in cell identity (Supplementary Fig.S1F). We next assessed endosomal processing by silencing Niemann–Pick type C1 (NPC1), a protein involved in cholesterol trafficking and endosomal maturation. NPC1 depletion significantly reduced luciferase expression (Fig. 1F), highlighting its importance in endosomal trafficking and mRNA release. Collectively, these results demonstrate that efficient delivery by L-36 LNPs depends on serum-derived protein adsorption, LDLR-mediated internalization, and NPC1-dependent endosomal processing, whereas other lipid-related receptors such as CD36 does not appear to play a major role in adipocyte uptake.

We next evaluated the membrane disruptive properties and endosomal escape behavior of L-36 LNPs to understand how they deliver mRNA efficiently while minimizing toxicity. We performed hemolysis assays at pH of 7.4 to assess the biocompatibility of L-36 LNPs. The results showed that L-36 caused less red blood cell lysis than Dlin-MC3 across ionizable lipid concentrations from 10 to 500 µM (Fig. 1G). This reduced hemolytic activity suggests that L-36 is more biocompatible and less disruptive to membranes under neutral conditions, which likely contributes to its improved safety profile. To see if L-36 could still promote endosomal escape without causing much membrane damage, we examined the activation of the galectin-8 pathway using Gal8–YFP–expressing cells^18^. In this assay, Gal8 puncta formation indicates localized endosomal rupture, which is necessary for cytosolic release of mRNA. Cells treated with L-36 LNPs showed a significantly higher number of Gal8 puncta compared to those treated with Dlin-MC3 LNPs (Fig. 1H-I). This suggests that L-36 facilitates efficient endosomal disruption at the intracellular level. The combination of reduced hemolytic activity at pH 7.4 and increased Gal8 recruitment within cells indicates that L-36 achieves a favorable balance between stability in extracellular environments and responsiveness to the acidic, cholesterol-rich endosomal environment. These data indicate that L-36 can trigger controlled endosomal release of mRNA without being overly disruptive to membranes at physiological pH. This provides a basis for designing ionizable lipids that combine high delivery efficiency with improved biocompatibility.

### L-36 LNPs enhance RNA delivery while minimizing inflammatory responses in human adipose and immune cell models

To evaluate the delivery performance and inflammatory profile of L-36 LNPs, we used a human iPSC-derived adipocyte–macrophage MPS that recapitulates key features of the subcutaneous adipose microenvironment and enables simultaneous assessment of mRNA delivery and inflammatory signaling ^19, 20^ (Fig. 2A). The MPS contains isogenic adipocytes and non-polarized macrophages at a physiological 1:10 ratio, which allows precise detection of both LNP uptake and cytokine induction^21, 22^. When LNPs with GFP mRNA were perfused through the MPS, L-36 produced significantly higher fluorescence intensity in adipocytes compared to Dlin-MC3, indicating enhanced mRNA delivery and reporter expression under human adipose tissue-like conditions (Fig. 2B). In addition, fluorescein-labeled control siRNA encapsulated in L-36 LNPs showed greater uptake than Lipofectamine, suggesting that L-36 also improves intracellular delivery of short RNA cargos (Fig. 2C)^23^. We next evaluated inflammatory signaling associated with LNP treatment. L-36 LNPs induced minimal changes in IL-6, TNFα, and CCL2 transcript levels whereas Dlin-MC3 triggered robust upregulation of these inflammatory markers (Fig. 2D–F). These findings suggest that L-36 has improved delivery activity and induces lower inflammatory signaling than Dlin-MC3 under the tested conditions. These results warrant further evaluation of L-36 as a candidate platform for adipose-directed RNA delivery. To further evaluate inflammatory responses in vivo, BALB/c mice were challenged with lipopolysaccharide (LPS)^24^ before administration of L-36 or Dlin-MC3 LNPs (Fig. 2G). Serum cytokine analysis showed lower IL-6 levels in mice treated with L-36 relative to Dlin-MC3 treated animals. PBS treated mice served as the controls for the study (Fig. 2H). To further assess inflammatory responses in primary human immune cells, peripheral blood mononuclear cells (PBMCs) were treated with L-36 or Dlin-MC3 LNP formulations under identical treatment conditions (Fig. 2I). Dlin-MC3 treatment resulted in significantly higher IL-6 production than L-36 (Fig. 2J), indicating that L-36 induces comparatively lower inflammatory signaling in primary human immune cells. These findings are consistent with the reduced inflammatory responses observed in the adipose MPS and in vivo cytokine studies, further supporting the use of L-36 as a platform for adipose-directed RNA delivery. Overall, these results demonstrate that L-36 LNPs support efficient RNA delivery in adipose tissue models while exhibiting reduced inflammatory responses relative to DLin-MC3 formulations in both ex vivo and in vivo settings, establishing L-36 as a promising and well-tolerated platform for adipose-directed RNA delivery and localized modulation of metabolic pathways.^25, 26^

**Fig. 2.**
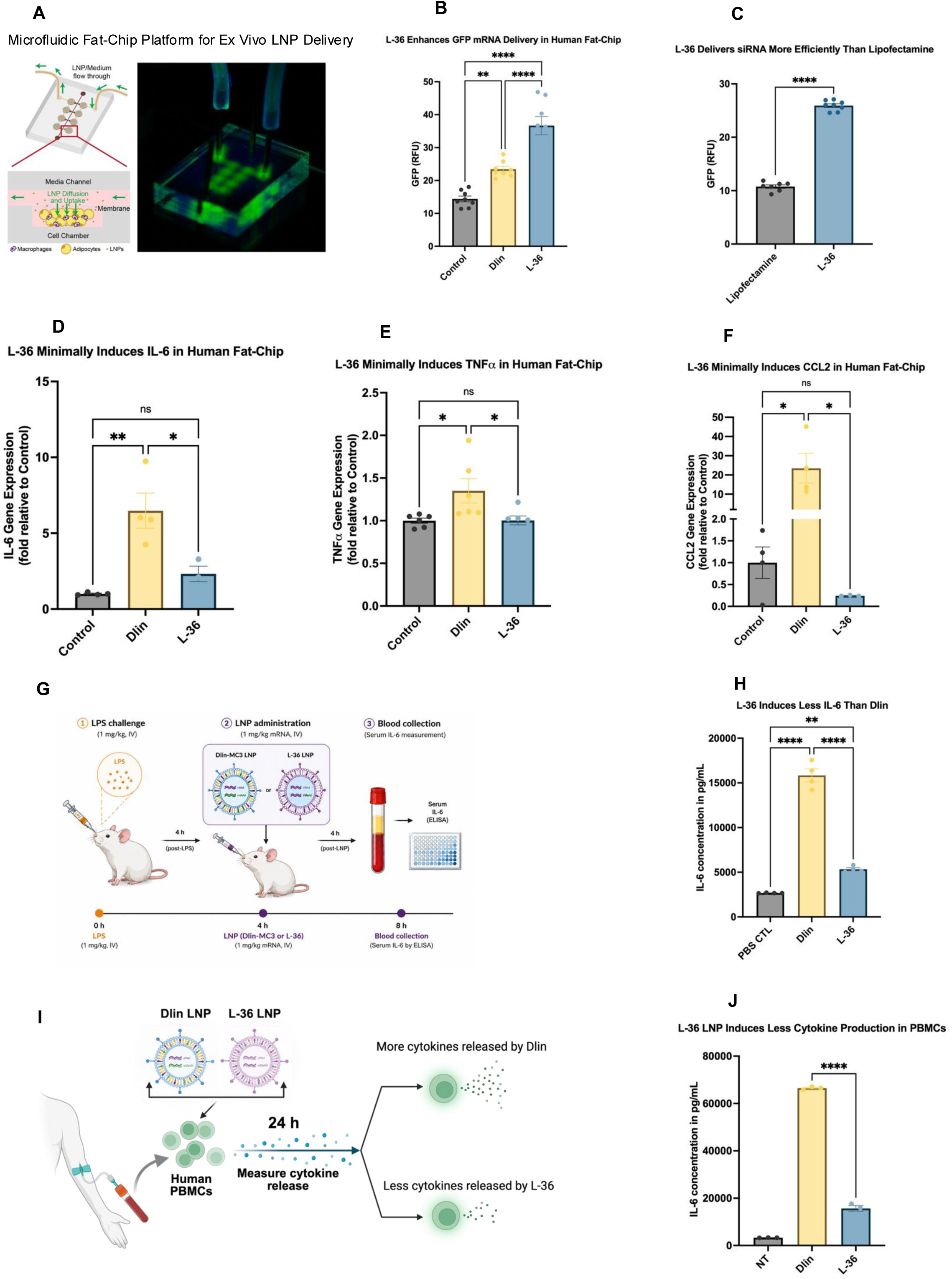
L-36 LNPs mediate efficient RNA delivery with reduced inflammatory activation in human adipose tissue models, PBMCs, and mice. (A) Schematic (left) and representative image (right) of the human adipose tissue microphysiological system (fat-chip) used to evaluate LNP-mediated RNA delivery and inflammatory responses. (B) GFP expression in adipocytes following delivery of GFP mRNA encapsulated in L-36 or DLin-MC3 LNPs within the fat-chip platform (n = 5–7). (C) Uptake of fluorescein-conjugated non-targeting control siRNA delivered using L-36 LNPs or Lipofectamine in human adipocytes (n = 7). (D–F) Relative expression of IL-6 (D), TNF-α (E), and CCL2 (F) in the adipose microphysiological system following treatment with L-36 or DLin-MC3 LNPs (n = 2–6). (G) Schematic of the acute inflammation model. BALB/c mice received intravenous lipopolysaccharide (LPS; 1 mg kg^−1^), followed 4 h later by intravenous administration of LNPs. Blood was collected 4 h after LNP administration for serum cytokine analysis. (H) Serum IL-6 concentrations measured by ELISA following LPS challenge and treatment with PBS, DLin-MC3 LNPs, or L-36 LNPs (n = 4 mice per group). (I) Schematic of the human PBMC assay used to evaluate inflammatory responses following LNP treatment. (J) IL-6 secretion from human peripheral blood mononuclear cells (PBMCs) following treatment with DLin-MC3 or L-36 LNP formulations (n = 3). Data are presented as mean ± SEM. Statistical significance was determined using one-way ANOVA followed by Tukey’s multiple-comparisons test. *P < 0.05, **P < 0.01, ***P < 0.001, ****P < 0.0001. Unless otherwise indicated, n represents independent biological replicates.

### BLNP-mediated silencing of TLE3 and ZFP423 promotes thermogenic gene expression in white adipocytes

To determine whether silencing transcriptional repressors of thermogenesis could enhance beige adipocyte-associated gene expression, we targeted TLE3 and ZFP423, two negative regulators of PRDM16 signaling (Fig. 3A). As illustrated in Fig. 3A, TLE3 interferes with the interaction between PRDM16 and PPARγ, whereas ZFP423 limits EBF2-mediated activation of the PRDM16 promoter^27, 28^. Because both pathways converge on PRDM16 activity, simultaneous suppression of TLE3 and ZFP423 was hypothesized to enhance thermogenic gene expression in white adipocytes.

**Fig. 3.**
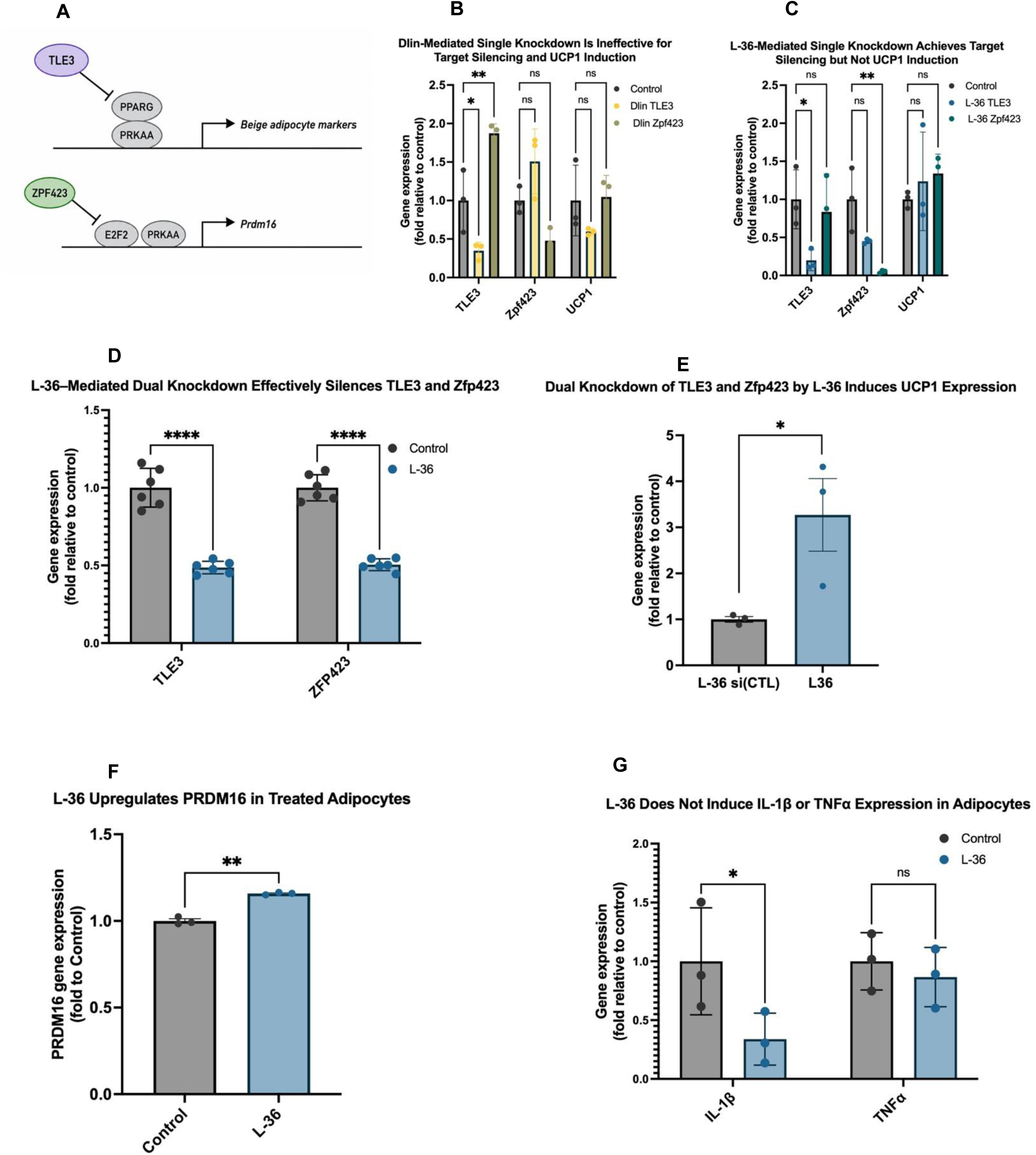
Dual silencing of TLE3 and ZFP423 by L-36 LNPs induces thermogenic gene expression in adipocytes without inflammatory activation. (A) Schematic illustrating the roles of TLE3 and ZFP423 in repressing PRDM16-dependent thermogenic gene expression in white adipocytes. TLE3 interferes with PRDM16–PPARγ signaling, whereas ZFP423 limits EBF2-mediated activation of the PRDM16 promoter. (B) SVF-derived white adipocytes treated with DLin-MC3 LNPs encapsulating siRNAs targeting TLE3 or ZFP423. DLin-MC3 efficiently suppressed TLE3 expression but exhibited limited activity against ZFP423 and did not induce UCP1 expression (n = 3). (C) SVF-derived adipocytes treated with L-36 LNPs encapsulating siRNAs targeting either TLE3 or ZFP423 exhibited efficient knockdown of the corresponding target gene; however, individual silencing of either gene was insufficient to induce UCP1 expression (n = 3). (D) Simultaneous delivery of siRNAs targeting TLE3 and ZFP423 using L-36 LNPs resulted in efficient dual-gene silencing (n = 6). (E) Dual knockdown of TLE3 and ZFP423 increased UCP1 transcript expression, whereas individual targeting of either gene alone did not induce UCP1 expression (n = 3). (F) PRDM16 expression was elevated following dual knockdown, consistent with activation of the thermogenic transcriptional program (n = 3). (G) Relative expression of IL-1β and TNF-α following treatment with L-36 LNPs containing siRNAs targeting TLE3 and ZFP423 (n = 3). Data are presented as mean ± SEM. Statistical significance was determined using one-way ANOVA for panels B–C and unpaired two-tailed Student’s t-tests for panels D–G. *P < 0.05, **P < 0.01, ****P < 0.0001. Unless otherwise indicated, n represents independent biological replicates.

We first optimized siRNA sequences targeting TLE3 and ZFP423 by screening a panel of Dicer-substrate siRNAs in white adipocytes differentiated from SVF of murine inguinal white adipose tissue.

From this screen, si(TLE3)#1 and si(ZFP423)#2 produced the strongest target knockdown and were selected for subsequent studies (Supplementary Fig.S1G). These optimized siRNAs were then subsequently encapsulated in L-36 LNPs to evaluate whether simultaneous silencing of TLE3 and ZFP423 could promote thermogenic gene expression in white adipocytes.

Adipocytes were treated with L-36 LNPs containing either a single siRNA (1.5 µg) or a combination of siTLE3 and siZFP423 (1.5 µg each; 3 µg total), and cells were collected after 48 hours for gene expression analysis. Dlin-MC3 formulation efficiently suppressed TLE3 but showed limited activity against ZFP423 (Fig. 3B). In contrast, L-36 LNPs produced substantial knockdown of both target transcripts following individual siRNA delivery (Fig. 3C). Although individual gene silencing was efficient, neither treatment significantly increased UCP1 expression (Fig. 3B,C), indicating that simultaneous inhibition of both transcriptional repressors is required to activate the thermogenic program. Co-encapsulation of siRNAs targeting both TLE3 and ZFP423 in L-36 LNPs resulted in dual gene suppression and increased UCP1 transcript levels by approximately 2.5-fold (Fig. 3D–E). PRDM16 expression was also elevated (Fig. 3F), consistent with enhanced thermogenic transcriptional signaling.

Dual siRNA treatment did not significantly alter IL-1β or TNF-α expression (Fig. 3G), indicating that thermogenic gene activation occurred without detectable inflammatory activation. These results identify simultaneous inhibition of TLE3 and ZFP423 as an effective strategy for activating thermogenic gene expression and provide the rationale for evaluating BLNPs in vivo.

### BLNPs mediate in vivo silencing of TLE3 and ZFP423 and promote thermogenic remodeling of iWAT

To evaluate the ability of L-36 LNPs to deliver RNA to adipose tissue in vivo, we first examined mRNA delivery to inguinal white adipose tissue (iWAT) in diet-induced obese mice. A schematic of the single-injection study design is shown in Fig. 4A. Bioluminescence imaging following administration of luciferase mRNA/LNP administration showed a strong localized signal within the injected adipose depots. Mice injected with L-36 LNPs exhibited intense bilateral luminescence in both iWAT depots, whereas PBS-treated controls showed no detectable signal (Fig. 4B–C). Analysis of excised adipose tissue further confirmed efficient local delivery of luciferase mRNA to the injected iWAT depots.

**Fig. 4.**
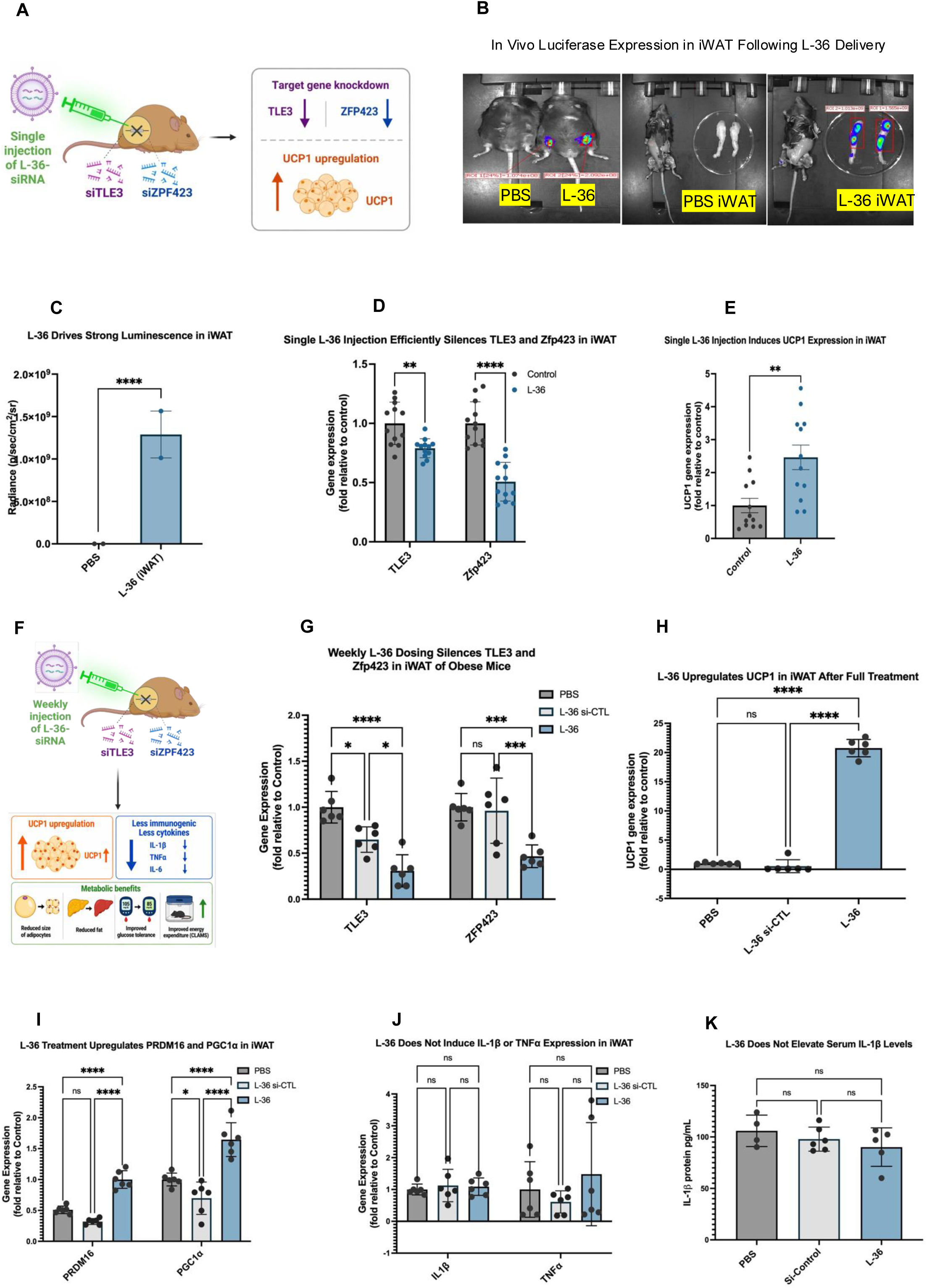
Single and repeated administration of L-36 LNPs induces and sustains thermogenic gene expression in iWAT without detectable inflammatory activation. (A) Schematic of the single-dose study in which L-36 LNPs encapsulating siRNAs targeting TLE3 and ZFP423 were administered by intra-iWAT injection to obese mice. (B) In vivo evaluation of L-36-mediated mRNA delivery following intra-iWAT administration. Left, IVIS imaging of mice injected with PBS or L-36 LNPs encapsulating luciferase mRNA. Middle, dissected iWAT from PBS-treated mice. Right, dissected iWAT from L-36-treated mice showing localized luciferase expression within both iWAT depots. (C) Quantification of luminescence intensity measured from the bilateral iWAT depots shown in panel B. (D) RT–qPCR analysis of iWAT collected 48 h after a single intra-iWAT injection demonstrated efficient knockdown of TLE3 and ZFP423 relative to PBS-treated controls (n = 6 biological replicates with 2 technical replicates each). (E) UCP1 transcript expression following a single intra-iWAT injection of L-36 LNPs encapsulating siRNAs targeting TLE3 and ZFP423 compared with PBS-treated controls (n = 6 biological replicates with 2 technical replicates each). (F) Experimental timeline of the repeated-dosing study. Mice received intra-iWAT injections on days 1, 7, 14, and 18, followed by molecular and metabolic analyses. (G) RT–qPCR analysis of TLE3 and ZFP423 expression in iWAT following repeated administration of PBS, scramble siRNA LNPs, or L-36 LNPs encapsulating siRNAs targeting TLE3 and ZFP423 (n = 6 mice per group). (H) UCP1 transcript expression in iWAT following repeated administration of PBS, scramble siRNA LNPs, or L-36 LNPs encapsulating siRNAs targeting TLE3 and ZFP423 (n = 6 mice per group). (I) Expression of thermogenic regulators PRDM16 and PGC1α in iWAT following repeated administration of PBS, scramble siRNA LNPs, or L-36 LNPs encapsulating siRNAs targeting TLE3 and ZFP423 (n = 6 mice per group). (J) Relative expression of IL-1β and TNF-α in iWAT following repeated administration of PBS, scramble siRNA LNPs, or L-36 LNPs encapsulating siRNAs targeting TLE3 and ZFP423 (n = 6 mice per group). (K) Serum IL-1β concentrations measured at the end of the repeated-dosing study in mice receiving PBS, scramble siRNA LNPs, or L-36 LNPs encapsulating siRNAs targeting TLE3 and ZFP423 (n = 6 mice per group). Data are presented as mean ± SEM. Statistical significance was determined using unpaired two-tailed Student’s t-tests. **P < 0.01, ***P < 0.001, ****P < 0.0001. Unless otherwise indicated, n represents independent biological replicates.

We next investigated whether L-36 mediated delivery of siRNAs targeting TLE3 and ZPF423 could promote thermogenic gene expression in vivo. Diet-induced obese mice maintained on a 60 % high-fat diet (Research Diets D12492) for 12 weeks were used for these studies. Animals received bilateral intra-WAT injections of L-36 LNPs encapsulating siRNAs targeting TLE3 and ZFP423 at a dose of 1.6 mg kg□^1^ siRNA, while PBS-treated mice served as controls. iWAT tissue collected 72 hours after injection showed a significant knockdown of both TLE3 and ZFP423 (Fig. 4D–E). This dual gene silencing was accompanied by increased UCP1 expression, consistent with enhanced thermogenic gene expression following inhibition of TLE3 and ZFP423 (Fig. 4E). These results demonstrate that BLNPs efficiently deliver siRNAs to adipose tissue in vivo, resulting in robust target gene silencing and initiation of thermogenic remodeling.

We next performed a repeated-dosing study to evaluate whether BLNP activity could be maintained during serial injection. Age-and weight-matched C57BL/6J male mice were fed a 60 % high-fat diet (HFD) for 8 weeks and then injected in their iWAT with 1.6 mg/kg of total TLE3 and ZFP423 siRNA per iWAT lobe 4 times over the course of 18 days (Fig. 4F). Mice injected with PBS or L-36 LNPs carrying scramble siRNA were used as controls. BLNPs treatment reduced TLE3 and ZFP423 mRNA levels in iWAT, with significant suppression of TLE3 observed relative to both PBS and scramble siRNA LNP controls (Fig. 4G). BLNPs administration increased UCP1 gene expression by approximately 20-fold compared with PBS and scramble siRNA treated animals (Fig. 4H). BLNP treatment increased PRDM16 and PGC1α expression, with PGC1α significantly elevated relative to scramble siRNA controls (Fig.4I). Together, these transcriptional changes are consistent with activation of the thermogenic signaling pathways following dual silencing of TLE3 and ZFP423.

Importantly, this transcriptional remodeling was not associated with detectable increases in inflammatory markers under the conditions tested. RT-qPCR analysis of inflammatory cytokines in iWAT showed no significant difference in IL-1β or TNF-α expression between BLNP-treated animals and either control group (Fig. 4J). Similarly, serum IL-1β levels remained comparable between BLNP-treated and control groups (Fig. 4K), suggesting limited systemic inflammatory activation under the conditions evaluated. Collectively, these findings demonstrate that repeated BLNP administration produces sustained target gene silencing and thermogenic remodeling of iWAT without detectable local or systemic inflammatory responses, supporting the evaluation of the metabolic benefits of localized adipose RNA therapy.

### BLNP treatment improves metabolic function in obese mice

To determine whether sustained thermogenic remodeling translates into metabolic improvements, we evaluated mice following the repeated BLNP dosing regimen shown in Fig. 4F. Repeated BLNP administration did not produce sustained differences in body weight compared with scramble siRNA controls, although a transient reduction was observed four days after the initial treatment (Fig. 5A). No significant difference in cumulative food intake was observed between BLNP-treated mice and control groups, indicating that treatment-associated metabolic changes were not accompanied by altered food consumption (Supplementary Fig. S2A). Despite minimal differences in body weight, BLNP-treated mice exhibited reduced fat mass compared with PBS controls (Supplementary Fig. S2B). Indirect calorimetry revealed significantly higher whole-body oxygen consumption (VO□) in BLNP-treated mice than in both PBS and scramble siRNA control groups during both the light and dark cycles (Fig. 5B,C). Similarly, carbon dioxide production (VCO□) was significantly elevated in BLNP-treated mice relative to both control groups during the light and dark phases (Fig. 5D). In contrast, respiratory exchange ratio (RER) values remained similar across all treatment groups during both light and dark cycles (Supplementary Fig. S2C), indicating no detectable changes in substrate utilization. In line with this, basal glucose levels were unchanged (Fig. 5E); however, serum glucose excursions following intraperitoneal glucose injections demonstrated improved glucose tolerance in BLNP-treated mice compared with scramble siRNA controls (Fig. 5E), particularly during the later phase of the glucose tolerance test, consistent with more efficient glucose clearance. Together, these findings demonstrate that repeated BLNP administration increased whole body respiration rates while improving glucose handling, consistent with enhanced whole-body metabolic function. To determine whether these metabolic adaptations were accompanied by structural remodeling of adipose tissue, we next examined iWAT morphology, thermogenic marker expression, and lipid metabolism.

**Fig. 5.**
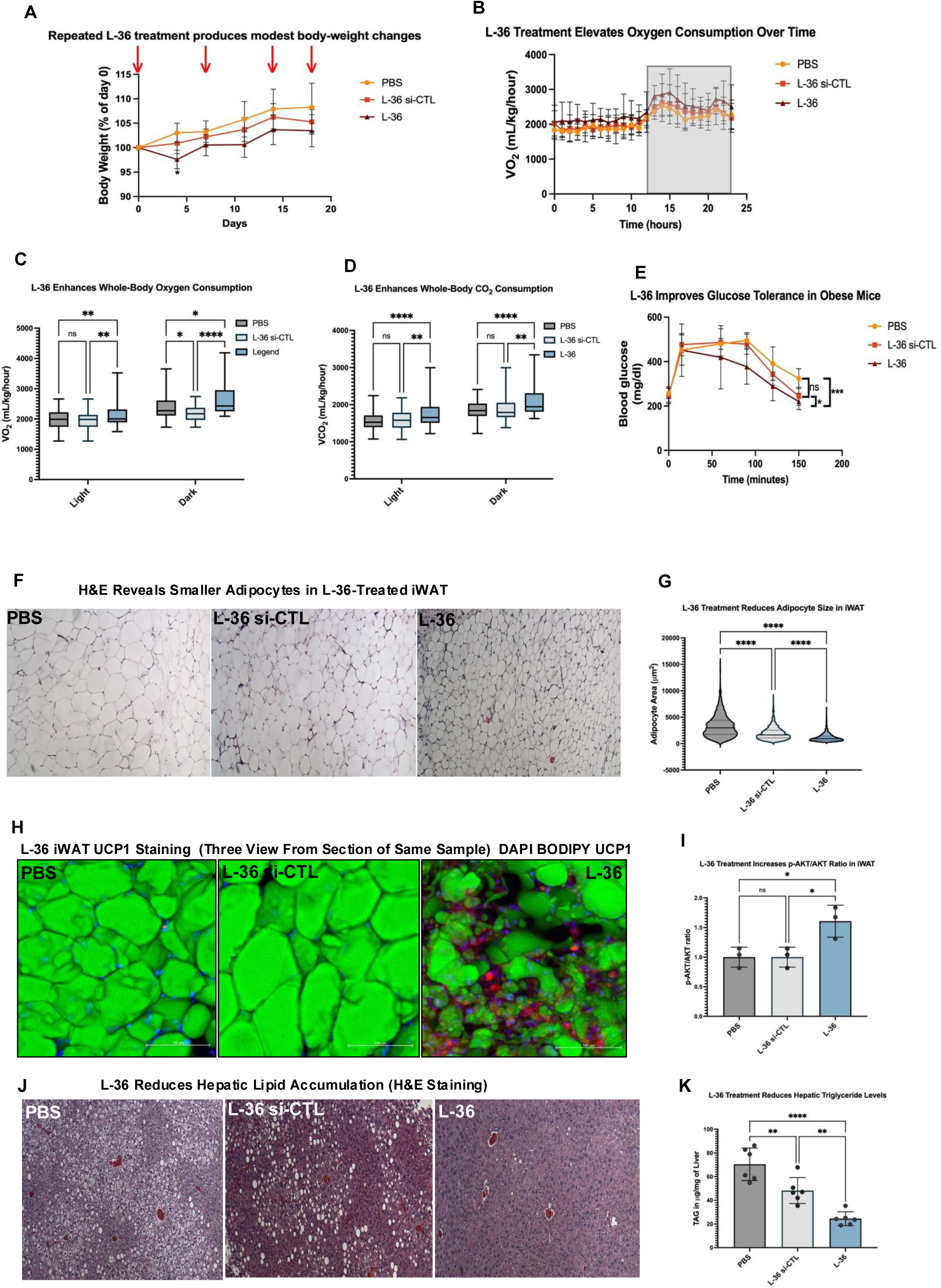
Repeated administration of L-36 LNPs targeting TLE3 and ZFP423 promotes adipose tissue remodeling and improves metabolic function in obese mice. (A) Body-weight trajectories of mice receiving PBS, scramble siRNA LNPs, or L-36 LNPs encapsulating siRNAs targeting TLE3 and ZFP423. Repeated L-36 administration produced modest changes in body weight while enabling substantial metabolic improvements. Statistical significance was determined using two-way repeated-measures ANOVA followed by Šídák’s multiple-comparisons test. (B) Whole-body oxygen consumption (VO₂) measured continuously by indirect calorimetry over the light and dark cycles. (C) Average whole-body oxygen consumption (VO₂) measured by indirect calorimetry. L-36-treated mice exhibited significantly higher oxygen consumption, indicating enhanced whole-body energy expenditure. (D) Whole-body carbon dioxide production (VCO₂) measured by indirect calorimetry. Increased VCO₂ further supports elevated metabolic activity following repeated L-36 treatment. (E) Glucose tolerance test (GTT) performed after repeated treatment. Mice were fasted for 6 h before intraperitoneal glucose administration (2.0 g kg^−1^ body weight). L-36-treated mice displayed improved glucose clearance, demonstrating enhanced glucose homeostasis. (F) Representative hematoxylin and eosin (H&E)-stained sections of inguinal white adipose tissue (iWAT). L-36 treatment markedly reduced adipocyte hypertrophy, consistent with adipose tissue remodeling. Scale bar, 100 μm. (G) Quantification of adipocyte area confirms a significant reduction in adipocyte size following repeated L-36 treatment. (H) Representative immunofluorescence images of iWAT stained for UCP1 (red), BODIPY (green), and DAPI (blue). Increased UCP1 expression demonstrates activation of a thermogenic adipocyte program following simultaneous silencing of TLE3 and ZFP423. (I) Quantification of phosphorylated AKT (Ser473) relative to total AKT (p-AKT/AKT). Enhanced AKT signaling indicates improved insulin-responsive signaling in adipose tissue after L-36 treatment. (J) Representative H&E-stained liver sections. L-36-treated mice exhibited reduced hepatic lipid accumulation, indicating improved systemic lipid metabolism. (K) Hepatic triglyceride (TAG) content was significantly reduced following repeated L-36 treatment, confirming attenuation of hepatic steatosis. Scramble siRNA LNP controls were included where indicated. Unless otherwise specified, n = 6 mice per group. For panels C and D, the center line represents the median and the box extends from the 25th to the 75th percentile. Data are presented as mean ± SEM. Statistical significance was determined using unpaired two-tailed Student’s *t*-test unless otherwise indicated. *P* < 0.05, P < 0.01, *P* < 0.001, P < 0.0001. Unless otherwise indicated, *n* represents independent biological replicates.

Hematoxylin and eosin (H&E) staining of the iWAT sections confirmed these changes at the cellular level as the adipocytes were smaller in the BLNP group compared with PBS and scramble siRNA controls (Fig. 5F). Quantitative analysis of the adipocyte area demonstrated a significant reduction in cell size following BLNP treatment (Fig. 5G). Consistent with activation of the thermogenic program, immunofluorescence staining demonstrated increased UCP1 expression throughout iWAT in BLNP-treated animals (Fig. 5H). Gross images of the injected iWAT depots are shown in Supplementary Fig. S3A. To determine whether these changes were associated with improved insulin signaling, we analyzed AKT activation in iWAT. Quantification of western blot analysis revealed a significantly increased p-AKT/AKT ratio in BLNP-treated mice compared with both control groups (Fig. 5I), whereas representative immunoblots are shown in Supplementary Fig. S3B.

Effects of BLNP treatment were also observed in the liver, where histological analysis revealed reduced lipid droplet accumulation in BLNP–treated mice relative to control groups (Fig. 5J). Quantification of hepatic triglyceride demonstrated significantly lower liver triglyceride content in BLNP-treated mice compared with both PBS and scramble siRNA controls (Fig. 5K). Serum triglyceride measurements similarly showed reduced circulating TAG levels in BLNP-treated mice relative to both controls (Supplementary Fig. S3C).

Together, these findings demonstrate that repeated BLNP administration promotes thermogenic remodeling of adipose tissue, improves adipose insulin signaling, and reduces hepatic lipid accumulation, indicating that localized adipose RNA therapy produces beneficial systemic metabolic effects.

## DISCUSSION

Adipocytes play a central role in metabolic homeostasis, and their dysfunction contributes to the development of metabolic disease. Thus, there is considerable interest in developing RNA delivery systems capable of efficiently delivering nucleic acids to adipocytes both in vitro and in vivo while maintaining acceptable tolerability profiles. However, efficient and well-tolerated platforms for RNA delivery to adipocytes remain limited. To date, only a limited number of delivery platforms have demonstrated efficient RNA delivery to adipocytes in vivo, including fluorinated polylysine-based nanoparticles^14, 29^. Yet, the translational potential of polylysine alkylated with fluorinated lipid tails is uncertain given the high toxicity of polylysine. Furthermore, therapeutic strategies requiring repeated RNA delivery to adipose tissue place particularly stringent demands on delivery efficiency and long-term biocompatibility. LNPs represent an attractive platform for adipose-directed RNA delivery; however, relatively few studies have systematically evaluated their performance in adipocytes or their suitability for repeated local administration. For example, Davies et al.^30^ demonstrated that subcutaneous injections of DLin-MC3 LNPs induced systemic increases in IL-6 and local injection-site inflammation, concluding that LNPs composed of DLin-MC3 lack the biocompatibility required for chronic subcutaneous administration. In addition, conventional cationic transfection reagents such as Lipofectamine exhibit poor transfection efficiency in adipocytes. Together, these findings highlight the need for RNA delivery platforms that combine efficient adipocyte transfection with the biocompatibility required for repeated local administration. However, systematic evaluation of structurally diverse ionizable lipids for RNA delivery to adipocytes has remained largely unexplored. Consequently, the structural features required for efficient adipocyte RNA delivery remain poorly understood.

In this report we developed a screening pipeline to identify ionizable LNPs capable of efficient RNA delivery to primary adipocytes using SVF-derived adipocytes as the discovery platform. This strategy successfully identified L-36 as a lead ionizable lipid that efficiently mediated adipocyte RNA delivery both in vitro and in vivo. Efficient adipocyte transfection was validated using both mRNA and siRNA cargoes across in vitro and in vivo models. L-36 LNPs were associated with lower inflammatory responses than DLin-MC3 formulations across multiple experimental models. L-36 has a unique structure for an ionizable lipid, it contains 3 lipid tails and also a double bond in each lipid tail. Incorporating a double bond into a lipid tail has been shown to enhance endosomal disruption and is likely playing a similar role here. LNPs made with L-36 have been previously investigated *in vivo* in both mice and NHPs after an intravenous injection, and it transfected the liver efficiently, with low toxicity, suggesting that it is generating a protein corona similar to traditional LNPs^17^. Mechanistic studies further indicated that L-36 uptake in adipocytes is largely mediated by LDLR, consistent with previous observations for conventional LNP formulations. Despite evidence that both formulations utilize LDLR-associated uptake pathways, the superior delivery performance of L-36 suggests that receptor engagement alone is insufficient to explain its activity in adipocytes. These findings point toward additional structural or intracellular trafficking features that may contribute to adipocyte RNA delivery. Interestingly, NPC1 knockdown inhibited L-36 transfection. Because NPC1 regulates cholesterol-dependent endosomal maturation, its involvement points to a distinct intracellular trafficking requirement for L-36. To our knowledge, NPC1 has not previously been implicated to LNP-mediated RNA delivery in adipocytes, and its involvement here suggests that L-36 depends on intracellular trafficking steps that are not yet fully defined, including mechanisms that ultimately enable it to reach the cytosol.

Similarly, our findings support the idea that thermogenic adipocytes are kept in a suppressed state by regulatory factors that limit their activation in normal conditions. D’Silva et al.^31^ recently identified several of these “thermogenic silencers,” such as ZFP423, HoxC10, IRF3, and MYPT1, which suppress thermogenic gene expression and limit the metabolic potential of adipocytes. ZFP423, in particular, acts as a major barrier to PRDM16 activity, which aligns with our observations. By delivering siRNAs targeting both TLE3 and ZFP423, BLNP treatment promoted thermogenic remodeling of white adipose tissue, as evidenced by robust UCP1 induction in both in vitro and in vivo models. These results support the thermogenic silencer model and show that targeted RNA delivery can modulate these pathways in a controlled and localized manner. Repeated BLNP administration produced metabolic improvements despite minimal sustained differences in body weight relative to scramble siRNA controls, indicating that the metabolic benefits are directly mediated by the browned adipose tissue rather than secondary to weight loss. Increased oxygen consumption, carbon dioxide production, reduced adipocyte size, enhanced UCP1 expression, improved AKT signaling, and reduced hepatic lipid accumulation together indicate that localized thermogenic remodeling can improve systemic metabolic function independently of substantial weight loss. Because scramble siRNA formulations also contain the L-36 delivery vehicle and were administered locally to adipose tissue, future studies will be needed to distinguish the relative contributions of RNA-mediated thermogenic remodeling and any siRNA-independent effects of the L-36 formulation on systemic metabolism.

Our findings demonstrate that BLNPs enable localized RNA delivery to adipose tissue resulting in sustained thermogenic remodeling in vivo. This platform may facilitate future mechanistic studies aimed at understanding how thermogenic pathways influence adipose tissue biology and systemic metabolism. For example, activation of BAT via cold exposure^32^ and implantation of genome engineered BeAT like adipocytes^26^ showed promising effects on reducing tumor growth in multiple oncological animals models presumably via competition for glucose between BAT and cancer cells^33^ fueled by the Warburg effect^34^. How much browning is required for this exciting effect, whether it can be applied to tumors predominantly utilizing fatty acids rather than glucose, such as certain intrahepatic cholangiocharinomas^35^, and how to translate it into clinical use remains to be resolved but could now be tested using BLNPs. Overall, BLNPs establish a biocompatible platform for localized adipose RNA delivery and demonstrate that sustained thermogenic remodeling can improve systemic metabolic function. These findings provide a foundation for developing adipose-targeted RNA therapeutics for metabolic disease and for investigating the broader physiological roles of thermogenic adipose tissue.

## CONCLUSIONS

In summary, we developed a biocompatible lipid nanoparticle system capable of delivering siRNAs to adipose tissue and triggering thermogenic remodeling *in vivo*. Using the three-tailed ionizable lipid L-36 to deliver siRNAs targeting TLE3 and ZFP423, we achieved localized thermogenic remodeling of white adipose tissue, characterized by efficient target gene silencing and increased expression of thermogenic markers. Repeated administration promoted sustained UCP1 expression, increased energy expenditure, improved glucose handling, reduced hepatic lipid accumulation, and minimal inflammatory responses under the conditions evaluated. These findings demonstrate that localized remodeling of a limited adipose tissue depot can produce measurable improvements in whole-body metabolic function. The comparatively low inflammatory responses observed with L-36 relative to conventional formulations support its further evaluation for repeated adipose-directed RNA delivery. Because this platform can be adapted to deliver siRNAs against other metabolic regulators, it may provide a versatile tool for investigating adipose tissue biology and metabolic regulation. Collectively, our findings establish BLNPs as a promising platform for localized adipose RNA delivery and provide a foundation for developing adipose-targeted RNA therapeutics to improve metabolic health.

## MATERIALS AND METHODS

### Isolation and Differentiation of Murine Adipocytes

SVF isolation: Stromal vascular fraction (SVF) cells were isolated from inguinal white adipose tissue (iWAT) depots of male C57BL/6J mice aged 4–8 weeks. Briefly, iWAT was minced and digested in Krebs-Ringer bicarbonate (KRB Buffer, 121mM NaCl, 4.9mM KCl, 1.2mM MgSO4•7H2O, 0.33mM CaCl2, 12mM HEPES, 3mM Glucose, pH 7.4) supplemented with 3% bovine serum albumin (BSA) and 0.1% type I collagenase. The digested tissue suspension was strained through a 100 μm and 40 μm cell strainer and centrifuged at 200 × g for 10 minutes. The resulting cell pellet was resuspended in erythrocyte lysis buffer, incubated for 5 min, and centrifuged again. Cells were resuspended in Dulbecco’s Modified Eagle Medium (DMEM) supplemented with 10% fetal bovine serum (FBS), 1 % penicillin and streptomycin (P/S). Culture media was additionally supplemented with 2.5 μg/mL amphotericin B during the first 3 days following isolation.

Adipogenic Differentiation: Preadipocytes isolated from the SVF of iWAT were cultured in DMEM supplemented with 10% FBS and 1% penicillin/streptomycin (Pen/Strep). Adipogenic differentiation was induced upon reaching 100% confluence using differentiation medium containing 0.2% Humulin N (Eli Lilly), 0.5 mM 3-isobutyl-1-methylxanthine (IBMX; Sigma-Aldrich), and 1 µM dexamethasone (Sigma-Aldrich). Three days after induction, cells were switched to maintenance media containing 10% FBS, 1% P/S, 0.2% Humulin N. On day six of differentiation, cells were switched to basal culture medium. For mRNA delivery experiment, cells were treated with LNPs containing luciferase mRNA on day 7-8 of differentiation. For serum-starvation experiments, cells were switched to FBS depleted medium on day 6 prior to LNP treatment. For chlorpromazine (CPZ) inhibition studies, cells were co-treated with 10 µM CPZ and LNPs on day 7-8 of differentiation. For siRNA delivery experiments, cells were seeded in 12-well plates and treated with LNPs formulations containing 1 µg siRNA per well on days 6-8 of differentiation. All siRNA sequences are provided in the Reagents and Tools table.

### Lipid Nanoparticle Formulation

Lipid stock solutions were prepared in ethanol at a concentration of 10 mg mL□^1^ using DLin-MC3-DMA (MedKoo, Cat. No. 555308), DOPE (Avanti Polar Lipids, Cat. No. 850725), cholesterol (Sigma-Aldrich, Cat. No. C8667), DMG-PEG2000 (Avanti Polar Lipids, Cat. No. 880151), and Lipid CL1 (L-36; Cayman Chemical, Cat. No. 38320). All stock solutions were stored at −30 °C until use. Prior to formulation, lipid stock solutions were thawed on ice and vortexed to ensure complete mixing. The cholesterol stock solution was briefly warmed to 40–50 °C to dissolve any crystallized material. DLin-MC3 LNPs were formulated at a molar ratio of DLin-MC3:DSPC:cholesterol = 50:10:38.5:1.5. L-series formulations were prepared using 34.41 mol% ionizable lipid, 24.72 mol% DOPE, 39.64 mol% cholesterol, and 1.22 mol% DMG-PEG2000. Complete formulation compositions are provided in Supplementary Table 1. Firefly luciferase (Fluc) mRNA or siRNA (1 μg μL□^−1^) was diluted in 25 mM citrate buffer (pH 4.0). The aqueous RNA solution was combined with the ethanol lipid phase at an aqueous-to-ethanol volume ratio of 1:3. Following mixing, formulations were briefly vortexed and incubated at room temperature for 10–15 min to allow nanoparticle self-assembly. Formulations were either used immediately or stored at 4 °C for short-term use. To remove residual ethanol and unencapsulated nucleic acids, LNP suspensions were dialyzed against a 100-fold excess volume of phosphate-buffered saline (PBS, pH 7.4) for 12 h at 4 °C using a Pur-A-Lyzer Mini 6000 dialysis device (Sigma-Aldrich, Cat. No. PURN60100-1KT).

### In Vitro LNP Screen Using a Luciferase Reporter Assay

SVF-derived adipocytes were seeded in 96-well plates and treated in biological triplicate with LNP formulations containing 333 ng firefly luciferase (Fluc) mRNA per well. Twenty-four hours after treatment, cells were lysed and luciferase expression was quantified using the Illumination™ Firefly Luciferase Enhanced Assay Kit (Gold Biotechnology) according to the manufacturer’s protocol. Luminescence was measured using a microplate reader and normalized as indicated in the corresponding figure legends.

### Dynamic light scattering (DLS) analysis of LNPs

Dynamic light scattering (DLS) was used to determine the hydrodynamic diameter and polydispersity index (PDI) of L-36 lipid nanoparticles. LNPs were prepared as described above. An aliquot of L-36 LNP suspension (20 μL) was diluted in phosphate-buffered saline (PBS, pH 7.4) to a final volume of 100 μL, corresponding to an approximate nucleic acid concentration of 0.05 mg mL□^−1^ where applicable. The diluted suspension was transferred to a disposable polystyrene cuvette (Sigma-Aldrich, Cat. No. 759200) and analyzed using a Zetasizer Nano ZS instrument (Malvern Panalytical) at 25 °C. Each sample was measured in triplicate, and the reported values represent the mean hydrodynamic diameter and PDI.

### Encapsulation Efficiency of L-36 LNPs

Encapsulation efficiency of L-36 LNPs was measured using a RiboGreen fluorescence assay, following the method described by Tanaka et al.^36^. LNP samples were incubated with RiboGreen in the presence or absence of 0.4% Triton X-100 to quantify total and unencapsulated mRNA. Fluorescence was read on a microplate reader, and encapsulation efficiency was determined by comparing mRNA signals from lysed and intact particles.

### siRNA knockdown of LDLR, CD36 and NPC1 expression

Predesigned siRNAs targeting mouse LDLR, CD36, and NPC1, as well as a non-targeting scramble control siRNA (MISSION siRNA; MilliporeSigma), were complexed with Lipofectamine RNAiMAX (Thermo Fisher Scientific) according to the manufacturer’s protocol. siRNA complexes were added to differentiating adipocytes on days 3 and 6 of differentiation at a dose of 20 pmol per well in 12-well plates or 1.6 pmol per well in 96-well plates. Cells were incubated overnight, after which the transfection medium was replaced with fresh culture medium. LNP treatment was performed on days 8–9 of differentiation.

### Hemolysis assay

Hemolysis was performed as previously reported with minor modifications^37^. Defibrinated red blood cells (RBCs; Hemostat Laboratories, Dixon, CA, USA) were washed in PBS (pH 7.4) by centrifugation at 750 × g for 10 min at 4 °C. The wash step was repeated three times and supernatant was removed after each cycle. Following the final wash, 200 µL of packed RBCs were diluted in 9.8 mL PBS (pH 7.4) to prepare a 2% (v/v) RBC suspension. LNP formulated with either L-36 or Dlin-MC3 were prepared at ionizable lipid concentrations ranging from 10 to 500 µM in PBS (pH 7.4). 100 µL aiquots of each formulation were incubated with 50 µL of the RBC suspension for 1 h at 37 °C. Samples were then centrifuged at 2,000 × g for 10 min, and hemoglobin release in the supernatant was quantified by measuring absorbance at 546 nm using a spectrophotometer. Complete hemolysis (100% control) was generated by incubating RBCs in deionized water. All experiments were performed in technical triplicate.

### Analysis of endosomal escape using Gal8–YFP–MDA-MB-231 cells

Gal8–YFP–expressing MDA-MB-231 cells were seeded in 24-well plates at a density of 1 × 10 cells per well and incubated overnight at 37 °C in DMEM supplemented with 10% FBS. LNPs formulated with Cy5-labeled mRNA were prepared using either L-36 or Dlin-MC3 as the ionizable lipid. The formulations were added to each well in 500 µL of fresh culture medium to achieve a final mRNA concentration of 200 µg mL□^−1^. Following a 4 h incubation, the treatment medium was removed, and cells were washed three times with PBS (pH 7.4). The cells were then fixed in 10% formalin at 4°C overnight. Cells were mounted using SlowFade™ Gold Antifade Mountant with DAPI (Invitrogen) prior to imaging. Fluorescence images were acquired and imaged with a Zeiss LSM880 confocal microscope. Gal8 puncta formation was used as a readout of endosomal membrane disruption and endosomal escape.

### Human WAT Chip Model

#### iPSC Cell Culture and Differentiation

All experiments used the healthy human male G15.AO iPSC line (RRID:CVCL_V192). All iPSCs were used at passage numbers ranging from 40 to 80. When iPSC colonies reached 80% confluence, they were dissociated using ReLeSR (#100-0483, Stemcell Technologies) and subcultured at a 1:10 ratio in mTeSR Plus medium (#100-0276, Stemcell Technologies) on Matrigel-coated substrates (#356231, Corning). Authentication was confirmed prior to experimentation by SNP analysis. Mycoplasma testing was conducted before all experiments and annually. Sterility was checked daily. The undifferentiated state, characterized by colony morphology, was verified before each differentiation. Pluripotency was assessed before all experiments and regularly, based on signature gene expression comparisons to endodermal and mesodermal differentiated states, and statistically analyzed using t-tests.

Differentiation of adipocytes were described in our previous study^20^. Briefly, all hiPSCs were firstly differentiated into mesenchymal progenitors (iPSC-MSCs) (STEMdiff Mesenchymal Progenitor Kit, #05240, Stemcell Technologies) and then transduced for chemically inducible PPARγ. After 48 hours of post-confluent culture, differentiation of iPSC-MSCs was induced in complete media (DMEM/F12 (#11320033, Gibco) containing 1% HEPES (#15630080, Gibco), 1% penicillin/streptomycin (#15140122, Gibco) and 10% fetal bovine serum (#EF-0500-A, Equafetal)) with supplements of 0.25 μM dexamethasone (#D1756, Sigma-Aldrich), 0.25 mM 3-isobutyl-1-methylxanthine (#I5879, Sigma-Aldrich), 100 nM rosiglitazone (#R2408, Sigma-Aldrich), and 500 nM insulin (#0002-8315-01, Humulin R, Eli Lilly) for 4 days (Day 0 to 3). Subsequent differentiation was completed in media supplemented with insulin and rosiglitazone at the same concentration (Day 4 to 14). Exogeneous PPARγ was induced by 1 μg/mL doxycycline (#D5207, Sigma-Aldrich) from Day 0 until the end. On Day 14, the iADIPOs expressed hallmark genes comparable to primary adipocytes collected from human biopsies^21^.

The M0 macrophages were generated using methods detailed in our previous publication{Qi, 2023 #7}. Briefly, hiPSCs were induced to hematopoietic stem cells (HSCs) using commercial kit (#05310, Stemcell Technologies). Floating HSCs were collected and further differentiated toward the human monocytes/macrophages in commercial media (#10961, Stemcell Technologies) with 100 ng/mL macrophage-colony stimulating factors (MCSF, #300-25-100UG, PeproTech, Thermofisher). We previously showed that these differentiated macrophages at nonactivated state were M2-like.

### MPS Culture and Coculture

MPS devices were fabricated in polydimethylsiloxane (PDMS, Sylgard 184, #NC9285739, FisherScientific) in accordance with our previous protocol^21^. Eight circular cell chamber units with diameter of 1500 μm and thickness of 60 μm per MPS were isolated from neighboring medium channel by polyethylene terephthalate (PET) isoporous membrane (#030060, TRAKETCH, SABEU GmbH & Co. KG) to ensure diffusive exchange between cells and medium.

The MPS was first loaded with differentiating iPSC-MSCs. Day 4 MSCs were dissociated (TrypLE Express, Gibco) and centrifuged as cell pellets. Cells were resuspended in triethanolamine buffer (TEOA; 0.3 M, pH 8) containing 3wt% adhesion peptide modified hyaluronic acid precursor and 0.3wt% MMP cleavable peptides (CQPQGLAKC, GenScript). The cell slurry with density of 8×10^7^ cells/mL was injected into the cell chamber and maintained for 30 min in incubator to remove uneven loading stress and allow for gelation. The MPS was connected to catheter couplers (#SC20/15 and #SP20/12, Instech Laboratories) and tubes (#06422-00, Cole-Parmer). The rest of adipocyte differentiation were in MPS with media supplied at a flow rate of 10 μL/hour using a syringe pump (#703007, Harvard Apparatus).

After Day 14, freely floating cells, which were presumed to be poorly differentiated, were flushed out by a continuous medium flow at a rate of 30 μL/min. Then dissociated macrophages at density of 5×10^6^ cells/mL were loaded into the cell chamber at flow rate of 5 μL/min. Loaded cell number was estimated based on cell loading density and loading volume. Coculture of two cells were in adipose medium with supply of 50 ng/mL M-CSF for 2 days to allow macrophage infiltration into adipocyte cluster.

### LNP Delivery and Characterization

After 2 days coculture, LNP solution after dialysis was diluted in 1 mL OptiMEM then flowed through MPS at a rate of 20 μL/hour. After 2 days LNP administration, MPS were rinsed by PBS, imaged, and snap frozen sequentially.

To compare delivery efficiency with conventional lipid–siRNA complexes, a fluorescein-conjugated control siRNA (#6201, Cell Signaling Technology) was diluted in 50 μL OptiMEM and combined with 50 μL Lipofectamine RNAiMAX (#13778150, Thermo Fisher). After 20 min incubation, the mixture was further diluted in OptiMEM to a final volume of 0.5 mL, delivering 10 pmol siRNA per MPS. In parallel, the same siRNA dose was encapsulated in LNPs and diluted in OptiMEM to the same final volume. Each formulation was perfused through monocultured WAT-MPS for 1 day at a rate of 20 μL/hour. Transfection efficiency was quantified by measuring epifluorescence intensity in each MPS well.

Brightfield and fluorescent images were captured by a wide-field fluorescence microscope (EVOS M5000 Imaging System, ThermoFisher). To enable fluorescent comparison, all staining and imaging procedures were kept consistent. ImageJ (no plug-ins) were used to quantify fluorescent intensity with consistent settings. Imaging analyses were based on whole cell chamber (under 4x objective lens, one chamber as one data point, 8 chambers per MPS).

Cells in MPS was collected by cutting and exfoliating the cell chamber off the MPS, and lysed in Trizol (#15596026, Invitrogen) for downstream gene expression analysis via RT-qPCR.

### LPS-Induced Systemic Inflammation Challenge

Wild-type BALB/c mice were used to evaluate inflammatory responses following LNP administration. Systemic inflammation was induced by intravenous (i.v.) administration of lipopolysaccharide (LPS; Sigma-Aldrich, St. Louis, MO, USA) at a dose of 1 mg kg□^−1^. The experimental design is illustrated in Figure 2G.

LNP formulations containing firefly luciferase (Fluc) mRNA were prepared and dialyzed as described above. Four hours following LPS challenge, mice received intravenous injections of L-36 or DLin-MC3 LNP formulations at a dose of 1.0 mg kg□^−1^ mRNA. PBS-treated mice served as controls.

Four hours after LNP administration, whole blood was collected and processed to obtain serum as described above. Serum interleukin-6 (IL-6) concentrations were quantified using a Mouse IL-6 ELISA Kit (Abcam, Cat. No. ab222503) according to the manufacturer’s instructions. Cytokine concentrations were calculated from standard curves generated using recombinant IL-6 standards.

### Human PBMC Cytokine Assay

Peripheral blood mononuclear cells (PBMCs) were isolated from fresh leukopaks obtained from healthy human donors (Stemcell Technologies, MA, USA). Following isolation, PBMCs were maintained in RPMI 1640 medium supplemented with 10% fetal bovine serum (FBS) and seeded into 96-well plates at a final volume of 100 μL per well. Cells were allowed to recover overnight at 37 °C in a humidified incubator containing 5% CO prior to treatment. LNP formulations containing firefly luciferase (Fluc) mRNA were prepared as described above using either L-36 or DLin-MC3 as the ionizable lipid. Formulations were diluted in Opti-MEM and added directly to PBMC cultures to achieve a final mRNA concentration of 1 μg mL□^1^. Cells were incubated with LNP formulations for 24 h at 37 °C. Following treatment, plates were centrifuged at 1000 × g for 10 min at 4 °C and cell culture supernatants were collected and stored at −80 °C until analysis. Interleukin-6 (IL-6) concentrations were quantified using a Human IL-6 ELISA Kit (Abcam) according to the manufacturer’s instructions. Absorbance was measured using a microplate reader and cytokine concentrations were determined from a standard curve generated using recombinant IL-6 standards. Data were collected from independent donor samples and analyzed to compare cytokine responses induced by L-36 and DLin-MC3 LNP formulations.

### In vivo LNP Administration

Intra-iWAT Administration: Mice were anesthetized with isoflurane delivered in oxygen. Anesthesia was induced using 3–4% isoflurane and maintained at 1–2% throughout the procedure. LNP formulations were prepared as described above and subsequently diluted in phosphate-buffered saline (PBS) to the desired RNA concentration prior to administration. For in vivo studies, formulations containing 10 μg firefly luciferase (Fluc) mRNA or 20 μg total siRNA were administered in a final injection volume of 75 μL per inguinal white adipose tissue (iWAT) depot. To achieve broad distribution throughout the adipose tissue, each iWAT depot received three separate injections of 25 μL administered at distinct locations within the tissue. Injections were performed near the surface of the depot to maximize local distribution while minimizing penetration into underlying tissues. For repeated dosing studies, diet-induced obese mice received bilateral intra-iWAT injections of PBS, L-36 LNPs containing scramble siRNA, or L-36 BLNPs containing siRNAs targeting TLE3 and ZFP423. Treatments were administered on days 1, 7, 14, and 18 of the study, and animals were monitored throughout the experimental period for subsequent metabolic and molecular analyses.

### Bioluminescence Imaging

Four hours after intra-iWAT administration of luciferase mRNA-loaded LNPs, mice received an intraperitoneal (i.p.) injection of D-luciferin sodium salt (150 μL, 15 mg mL□^1^). Ten minutes following D-luciferin administration, animals were anesthetized with isoflurane and imaged using an IVIS Xenogen bioluminescence imaging system (PerkinElmer). Bioluminescent images were acquired using standardized instrument settings and analyzed using Living Image software (PerkinElmer). Regions of interest (ROIs) were drawn around the inguinal white adipose tissue (iWAT) depots, and total photon flux (photons s ^1^) was calculated for each animal to quantify luciferase expression.

### Indirect Calorimetry

Whole-body metabolic parameters were assessed using a Comprehensive Lab Animal Monitoring System (CLAMS; Columbus Instruments). Following four intra-iWAT administrations of BLNPs containing siRNAs targeting TLE3 and ZFP423 over an 18-day treatment period, mice were individually housed in metabolic chambers maintained at 23 °C. Animals were acclimated to the chambers for 24 h prior to data collection. Oxygen consumption (VO□), carbon dioxide production (VCO□), and respiratory exchange ratio (RER) were continuously monitored according to the manufacturer’s protocols. Metabolic data were collected during both light and dark cycles and analyzed using CalR, a web-based analysis platform for indirect calorimetry experiments^38^.

### Glucose Tolerance Test

Glucose tolerance tests (GTTs) were performed following a 6 h fast with ad libitum access to water. Baseline blood glucose levels were measured from tail vein blood using a handheld glucometer and compatible glucose test strips. Mice were then administered D-glucose (2.0 g kg□^1^ body weight) by intraperitoneal (i.p.) injection. Blood glucose concentrations were measured at 15, 30, 60, 90, 120, and 150 min following glucose administration. Glucose excursion curves were generated from the resulting measurements, and the area under the curve (AUC) was calculated to assess glucose tolerance.

### Body Composition Analysis

Body composition was measured following four administrations of BLNPs containing siRNAs targeting TLE3 and ZFP423 over an 18-day treatment period. Fat mass and lean mass were quantified using an EchoMRI body composition analyzer (EchoMRI LLC, Houston, TX, USA) according to the manufacturer’s instructions. Briefly, mice were placed in a restraining tube and inserted into the measurement chamber for non-invasive analysis. Measurements were completed within 3–6 min per animal. Fat and lean mass were measured before the first BLNP administration and following completion of the treatment regimen. Body composition values were normalized to baseline measurements, and fat and lean mass were expressed as a percentage of total body weight.

### H&E Staining and Adipocyte Size Quantification

Inguinal white adipose tissue (iWAT) was harvested from mice treated with PBS, scramble siRNA LNPs, or BLNPs containing siRNAs targeting TLE3 and ZFP423. Tissues were fixed in 10% neutral-buffered formalin, paraffin embedded, sectioned, and stained with hematoxylin and eosin (H&E) according to standard histological procedures. Stained sections were imaged using an EVOS M5000 imaging system (Thermo Fisher Scientific).

Adipocyte morphology was quantified using the Adiposoft plugin implemented in Fiji/ImageJ^38^. Adipocyte cross-sectional area and diameter were measured from representative iWAT sections. One iWAT image from six mice were analyzed per group. The resulting measurements were used for quantitative comparison of adipocyte size between experimental groups.

### UCP1 Immunofluorescence Analysis of iWAT

Immunofluorescence staining of inguinal white adipose tissue (iWAT) was performed as previously described with minor modifications^39^. Briefly, iWAT depots were harvested and fixed in 4% paraformaldehyde overnight at 4 °C. Tissues were cryoprotected by sequential incubation in 15% and 30% sucrose solutions at 4 °C and subsequently embedded in Neg-50 mounting medium. Frozen tissue blocks were sectioned using a Leica CM3050S cryostat maintained at −35 °C.

Prior to staining, tissue sections were washed with phosphate-buffered saline (PBS) and blocked for 1 h at room temperature in blocking buffer consisting of HBSS supplemented with 10% fetal calf serum (FCS), 0.1% bovine serum albumin (BSA), 0.05% saponin, and 2% donkey serum. Sections were then incubated overnight at 4 °C with anti-UCP1 primary antibody (1:100 dilution; Cell Signaling Technology) prepared in blocking buffer.

The following day, sections were washed three times with blocking buffer and incubated for 2 h at room temperature with Alexa Fluor 594-conjugated donkey anti-rabbit IgG secondary antibody (1:800; Jackson ImmunoResearch) and BODIPY 493/503 (3 μM) to visualize neutral lipid droplets. Following staining, sections were mounted using SlowFade™ Gold Antifade Mountant containing DAPI (Invitrogen) and imaged using a Zeiss LSM880 confocal microscope.

### RNA Isolation and Quantitative RT-PCR

Total RNA was isolated from cultured cells or tissue samples using TRIzol Reagent (Invitrogen) and purified using the Monarch Total RNA Miniprep Kit (New England Biolabs, Cat. No. T2010S) according to the manufacturer’s instructions, including on-column DNase treatment. RNA concentration and purity were determined prior to downstream analysis.

Complementary DNA (cDNA) was synthesized using the Maxima First Strand cDNA Synthesis Kit (Thermo Fisher Scientific, Cat. No. K1672). Quantitative real-time PCR (RT-qPCR) was performed on a QuantStudio 5 Real-Time PCR System (Applied Biosystems) using TaqMan Universal Master Mix II and validated PrimeTime qPCR primer-probe assays (Integrated DNA Technologies). For each reaction, 10–20 ng of cDNA was used as template.

Relative gene expression levels were calculated using the comparative ΔΔCt method and normalized to the appropriate housekeeping gene. Primer and probe information are provided in the Reagents and Tools Table.

### Western Blot Analysis

Protein lysates were prepared from inguinal white adipose tissue (iWAT) using RIPA buffer (50 mM Tris-HCl, pH 8.0, 150 mM NaCl, 1% Triton X-100, 0.5% sodium deoxycholate, and 0.2% SDS) supplemented with Halt™ Protease and Phosphatase Inhibitor Cocktail (Thermo Fisher Scientific). Protein samples were mixed with Laemmli sample buffer and denatured at 95 °C for 5 min prior to analysis.

Equal amounts of protein (30 μg per lane) were separated on 4–20% Mini-PROTEAN TGX gradient gels (Bio-Rad) and transferred to nitrocellulose membranes using a Trans-Blot Turbo Transfer System (Bio-Rad). Membranes were blocked in 5% non-fat milk prepared in Tris-buffered saline containing 0.1% Tween-20 (TBST) and incubated overnight at 4 °C with primary antibodies diluted in blocking buffer.

Following primary antibody incubation, membranes were washed three times with TBST and incubated with IRDye 680LT goat anti-mouse IgG or IRDye 800CW goat anti-rabbit IgG secondary antibodies (LI-COR Biosciences) for 1 h at room temperature. Membranes were subsequently washed three times with TBST and imaged using an Odyssey Imaging System (LI-COR Biosciences). Band intensities were quantified using Image Studio Lite software. For membranes probed for multiple targets, primary and secondary antibody incubations were performed sequentially for each protein. Phosphorylated AKT levels were normalized to total AKT levels prior to statistical analysis. A complete list of antibodies is provided in the Reagents and Tools Table.

### IL-6 and IL-1β ELISA assay

Serum concentrations of interleukin-1β (IL-1β) and interleukin-6 (IL-6) were quantified using commercial ELISA kits (Mouse IL-1β ELISA Kit, ab197742, Abcam; Mouse IL-6 ELISA Kit, ab222503, Abcam) following the manufacturers’ instructions. Whole blood was collected from mice and allowed to clot undisturbed at room temperature for 15–30 minutes. The samples were then centrifuged at 2,000 × g for 10 minutes at 4 °C to separate the serum. The supernatant was carefully transferred into clean tubes and maintained at 2–8 °C during handling. Serum samples were stored at –20 °C until analysis to prevent degradation from repeated freeze thaw cycles.

For ELISA, serum samples and standards were added to antibody-coated wells and incubated according to the kit protocol. After sequential washing and incubation with horseradish peroxidase (HRP)–conjugated secondary antibodies, color development was achieved using a chromogenic substrate. The optical density was measured using a microplate reader, and cytokine concentrations were calculated from standard curves generated with known concentrations of recombinant IL-1β and IL-6.

### Triglyceride Quantification in Serum and Liver Tissue

Serum triglyceride (TAG) concentrations were measured using a Triglyceride Assay Kit (Abcam, Cat. No. ab65336) according to the manufacturer’s instructions. Serum samples were prepared as described above for cytokine analysis. TAG concentrations were determined from a standard curve generated using the supplied triglyceride standards and reported as mg dL□^1^.

Hepatic triglyceride content was quantified using the same assay following lipid extraction from liver tissue. Briefly, 40–50 mg of liver tissue was homogenized in 0.5 mL phosphate-buffered saline (PBS). Lipids were extracted by adding 1.6 mL chloroform:methanol (2:1, v/v), followed by centrifugation at 3,000 rpm for 10 min at room temperature to separate the organic and aqueous phases. The lower organic phase was collected and air-dried overnight in a chemical fume hood. Dried lipid extracts were resuspended in 800 μL of 1% Triton X-100 in absolute ethanol and vortexed until fully solubilized.

Triglyceride concentrations were measured using the Triglyceride Assay Kit according to the manufacturer’s protocol. Hepatic triglyceride content was calculated from a triglyceride standard curve, normalized to tissue weight, and reported as μg TAG mg□^1^ liver tissue.

### Animal Models

All animal procedures were approved by the University of California, Berkeley Institutional Animal Care and Use Committee (IACUC) and conducted in accordance with institutional guidelines and ethical regulations for animal research. Mice were housed under standard laboratory conditions at 23°C with ad libitum access to food and water and maintained on a regular chow diet (LabDiet 5053) unless otherwise indicated.

For obesity studies, age-matched male C57BL/6J mice were fed a 60% high-fat diet (Research Diets, D12492) for 12 weeks to induce obesity prior to single dose administration experiments and for 8 weeks prior to longitudinal, multiple dose experiments. Body weight was monitored twice weekly throughout the study. Additional details regarding treatment schedules and experimental endpoints are provided in the corresponding sections below.

### Ethical Statement

All studies were conducted in accordance with applicable institutional guidelines and regulations. Animal experiments were approved by the University of California, Berkeley Institutional Animal Care and Use Committee (IACUC). Studies involving human induced pluripotent stem cell (iPSC)-derived tissues were conducted under protocols approved by the Stem Cell Research Oversight (SCRO) Office at the University of California, Berkeley. Research activities involving recombinant nucleic acids and biological materials were performed under protocols approved by the Committee on Laboratory and Environmental Biosafety (CLEB) and the Office of Environment, Health & Safety (EH&S) at the University of California, Berkeley.

### Statistical Analysis

All experiments were performed using independent biological samples unless otherwise indicated. Data are presented as mean ± standard error of the mean (SEM). Statistical analyses were performed using GraphPad Prism software (GraphPad Software, San Diego, CA, USA).

For comparisons between two groups, statistical significance was determined using an unpaired two-tailed Student’s t-test. For experiments involving three or more groups, statistical significance was determined using one-way analysis of variance (ANOVA) followed by Tukey’s multiple-comparisons test. The statistical test used for each experiment is indicated in the corresponding figure legend. Differences were considered statistically significant at P < 0.05. Statistical significance is denoted as P < 0.05 (*), P < 0.01 (**), P < 0.001 (***), and P < 0.0001 (****).

### Reagents and Tools Table

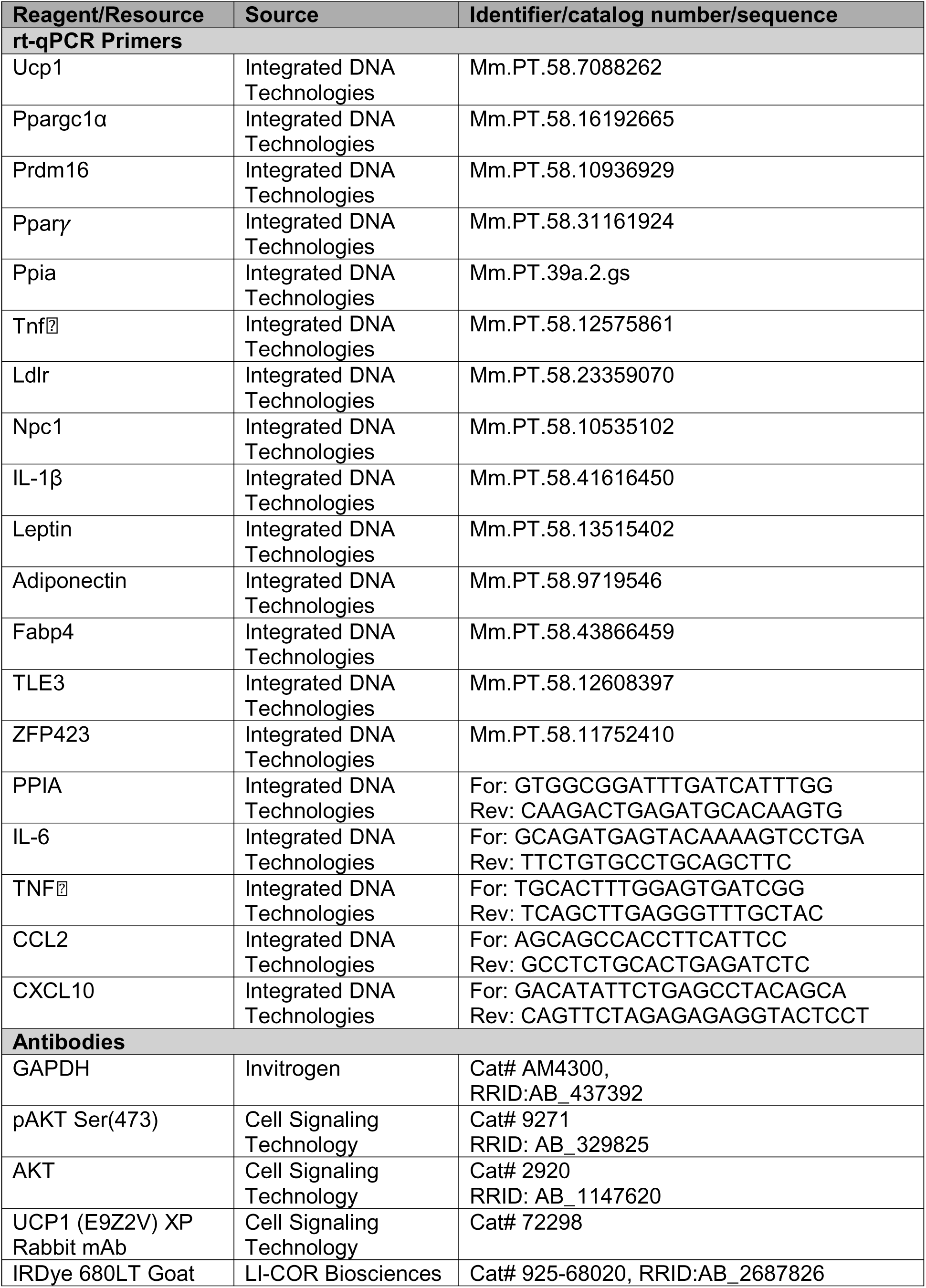
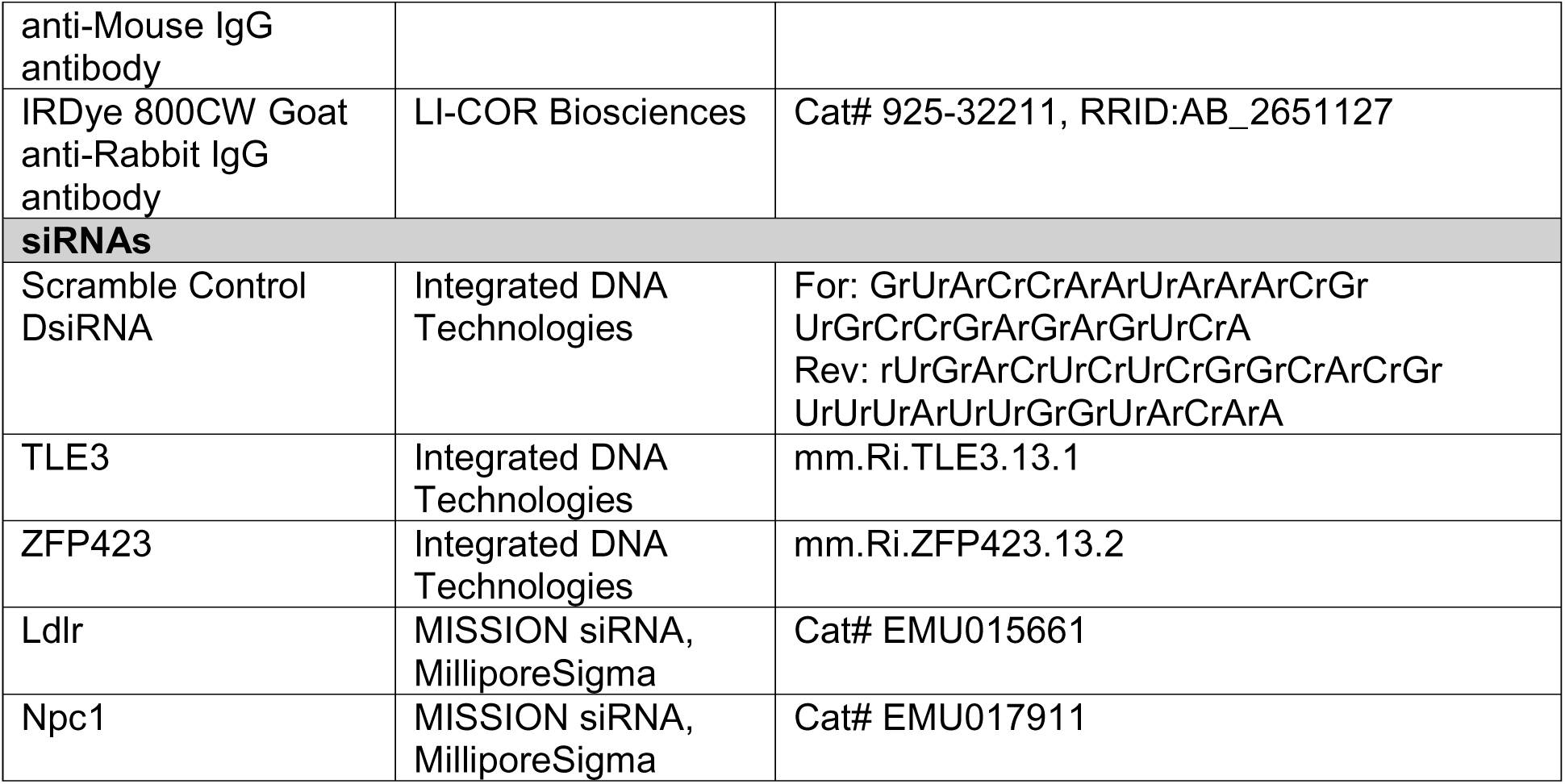

## Author contributions

RS, ALG, NM, and AS conceived and designed the study. RS and ALG contributed equally to the experimental work as joint first authors, with RS preparing the initial manuscript draft. RS, ALG, YH, LQ, NS, JL, NK, and VT performed the experiments, with LQ leading work related to the MPS system. RS, ALG, and YH analyzed and interpreted the data. NM and AS provided overall supervision, project direction, and critical input on the study design and interpretation of results. All authors contributed to manuscript revision and approved the final version.

## Data and materials availability

All data are available in the main text or the supplementary materials.

## Competing interests

The authors declare the following competing interests: N.M. own equity in Opus Biosciences. All the other authors declare no competing interests.

