## Supplementary Information for "Browning Lipid Nanoparticles Deliver Metabolic Benefits via Transient Conversion of White Adipose Tissue"

Rohit Sharma^1,2,†^ Amanda L Gunawan^3^,^†^ Yuchen He^3^, Lin Qi^3^, Nehal Singhal^1,2^, Jesslyn Lukman^3^, Nayiri Kalindjian^3^, Vyvylyn Tran^3^, Niren Murthy^1,2,^
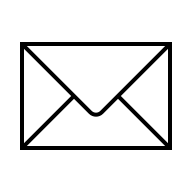
, Andreas Stahl^3,^
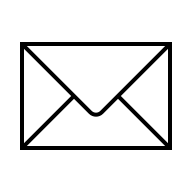

1 Department of Bioengineering, University of California, Berkeley, California 94720, United States

2 The Innovative Genomics Institute, 2151 Berkeley Way, Berkeley, California 94704, United States

3 Department of Nutritional Sciences and Toxicology, University of California Berkeley, California 94720, United States

† These authors contributed equally: Rohit Sharma and Amanda L Gunawan

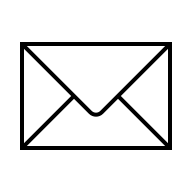

**Supplementary Table S1**

Composition of LNPs screened in this study. Formulations were prepared at a 34.41:24.72:0:39.64:1.22 molar ratio (ionizable lipid : DOPE : DSPC : cholesterol : DMG-PEG), unless otherwise indicated.

| LNP ID | Ionizable lipid 1 | Ionizable lipid 2 | DOPE | DSPC | Cholesterol | DMG-PEG |
| --- | --- | --- | --- | --- | --- | --- |
| L-36 | Lipid CL1 (34.41%) | 0 | 24.72% | 0 | 39.64% | 1.22% |
| L-32 | Lipid-2-2(8,8)-4C-CH₃ (34.41%) | 0 | 24.72% | 0 | 39.64% | 1.22% |
| Dlin | Dlin-MC3-DMA (50%) | 0 | 0 | 10% | 38.50% | 1.50% |
| L-24 | OF-02 (34.41%) | 0 | 24.72% | 0 | 39.64% | 1.22% |
| L-47 | Lipid III-45 (34.41%) | 0 | 24.72% | 0 | 39.64% | 1.22% |
| L-42 | ALC-0315 analog-1 (34.41%) | 0 | 24.72% | 0 | 39.64% | 1.22% |
| L-10 | CKK-E12 (34.41%) | 0 | 24.72% | 0 | 39.64% | 1.22% |
| L-5 | L-319 (34.41%) | 0 | 24.72% | 0 | 39.64% | 1.22% |
| L-46 | CL4F8-6 (34.41%) | 0 | 24.72% | 0 | 39.64% | 1.22% |
| L-45 | AA-T3A-C12 (34.41%) | 0 | 24.72% | 0 | 39.64% | 1.22% |
| L-4 | Lipid-5 (34.41%) | 0 | 24.72% | 0 | 39.64% | 1.22% |
| L-64 | L202 (34.41%) | 0 | 24.72% | 0 | 39.64% | 1.22% |
| L-8 | 306Oi10 (34.41%) | 0 | 24.72% | 0 | 39.64% | 1.22% |
| L-25 | Lipid A9 (34.41%) | 0 | 24.72% | 0 | 39.64% | 1.22% |
| L-2 | SM-102 (34.41%) | 0 | 24.72% | 0 | 39.64% | 1.22% |
| L-18 | ALC-0315 (34.41%) | 0 | 24.72% | 0 | 39.64% | 1.22% |
| L-1 | Dlin-MC3-DMA (30.78%) | GL67 (N⁴-cholesteryl-spermine·3HCl, 15%) | 20.79% | 0 | 32.17% | 1.23% |
| L-14 | CKK-E12 (30.74%) | GL67 (N⁴-cholesteryl-spermine·3HCl, 15%) | 20.84% | 0 | 32.15% | 1.25% |
| L-6 | Lipid A6 (34.41%) | 0 | 24.72% | 0 | 39.64% | 1.22% |
| L-40 | YK-009 (34.41%) | 0 | 24.72% | 0 | 39.64% | 1.22% |
| L-56 | C13-112 tetra-tail (34.41%) | 0 | 24.72% | 0 | 39.64% | 1.22% |
| L-15 | Lipid A9 (30.74%) | GL67 (N⁴-cholesteryl-spermine·3HCl, 15%) | 20.84% | 0 | 32.15% | 1.25% |
| L-7 | Lipid-5 (30.74%) | GL67 (N⁴-cholesteryl-spermine·3HCl, 15%) | 20.84% | 0 | 32.15% | 1.25% |
| L-66 | Lipid-14 (34.41%) | 0 | 24.72% | 0 | 39.64% | 1.22% |
| L-12 | 306Oi10 (30.74%) | GL67 (N⁴-cholesteryl-spermine·3HCl, 15%) | 20.84% | 0 | 32.15% | 1.25% |
| L-9 | C12-200 (34.41%) | 0 | 24.72% | 0 | 39.64% | 1.22% |
| L-16 | C13-112 tetra-tail (30.74%) | GL67 (N⁴-cholesteryl-spermine·3HCl, 15%) | 20.84% | 0 | 32.15% | 1.25% |
| L-11 | Lipid A6 (30.74%) | GL67 (N⁴-cholesteryl-spermine·3HCl, 15%) | 20.84% | 0 | 32.15% | 1.25% |
| L-3 | L-319 (30.74%) | GL67 (N⁴-cholesteryl-spermine·3HCl, 15%) | 20.84% | 0 | 32.15% | 1.25% |
| L-13 | C12-200 (30.74%) | GL67 (N⁴-cholesteryl-spermine·3HCl, 15%) | 20.84% | 0 | 32.15% | 1.25% |
| L-17 | Lipid-14 (30.74%) | GL67 (N⁴-cholesteryl-spermine·3HCl, 15%) | 20.84% | 0 | 32.15% | 1.25% |

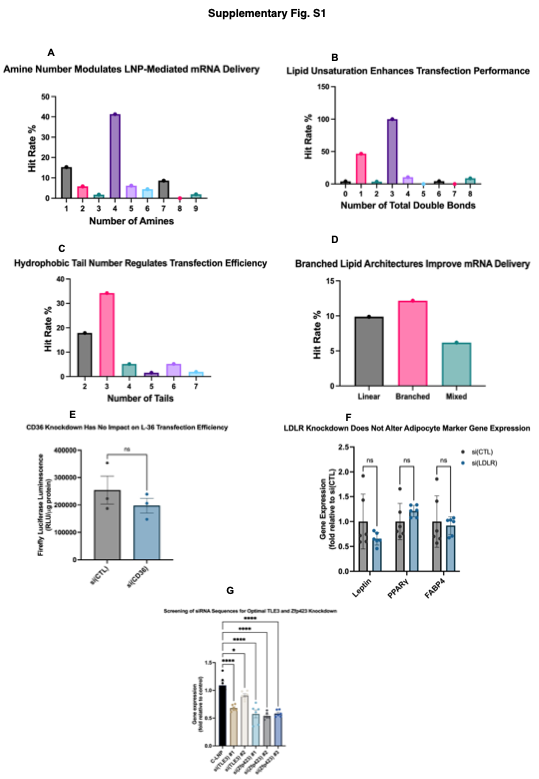

**Supplementary Fig. S1 | Structure-activity relationship analysis identifies key lipid structural features of L-36 and validates experimental controls for mechanistic studies.**

(A) Structure-activity relationship (SAR) analysis showing the effect of the number of amine groups on LNP-mediated mRNA delivery. Lipids containing four amine groups exhibited the highest transfection efficiency, demonstrating that amine number is a critical determinant of LNP performance. (B) Effect of lipid unsaturation on mRNA delivery. LNPs containing lipids with three total double bonds achieved the highest transfection efficiency, indicating that lipid unsaturation strongly influences intracellular delivery. (C) Effect of hydrophobic tail number on transfection efficiency. Lipids containing three hydrophobic tails outperformed alternative architectures, demonstrating the importance of tail number for efficient mRNA delivery. (D) Effect of lipid architecture on transfection efficiency. Branched lipid structures exhibited higher transfection efficiency than linear or mixed architectures, supporting branched lipid design for optimized intracellular delivery. (E) CD36 knockdown in differentiated adipocytes did not alter luciferase expression following L-36 LNP treatment, indicating that CD36 is not required for L-36-mediated cellular uptake or transfection. (F) LDLR knockdown did not significantly alter the expression of adipocyte marker genes, including Leptin, PPARγ, and FABP4, confirming that LDLR silencing does not disrupt adipocyte identity. (G) Screening of Dicer-substrate siRNA sequences targeting TLE3 and Zfp423 in SVF-derived white adipocytes. Transcript levels measured 48 h after treatment identified the most effective siRNA sequences, which were selected for all subsequent in vitro and in vivo studies.

Data are presented as mean ± SEM. Statistical significance was determined using unpaired two-tailed Student's *t*-test or one-way ANOVA with appropriate multiple-comparison correction, as described in the Methods. *P* < 0.05, *****P* < 0.0001; ns, not significant. Unless otherwise indicated, *n* represents independent biological replicates.

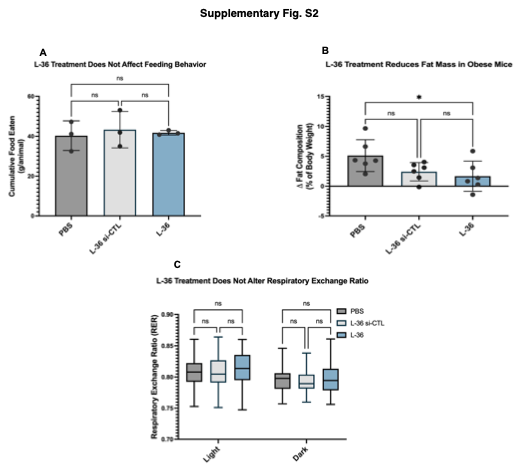

**Supplementary Fig. S2 | Additional metabolic characterization following repeated administration of L-36 lipid nanoparticles targeting TLE3 and ZFP423 in obese mice.**

(A) Cumulative food intake measured throughout the repeated treatment study. Food consumption was comparable between mice receiving PBS or L-36 LNPs encapsulating siRNAs targeting TLE3 and ZFP423, indicating that metabolic improvements were independent of changes in food intake (n = 3 cages per group). (B) Fat mass normalized to pre-treatment baseline following repeated administration of PBS, scramble siRNA LNPs (L-36 si-CTL), or L-36 LNPs encapsulating siRNAs targeting TLE3 and ZFP423. (C) Respiratory exchange ratio (RER) measured by indirect calorimetry, demonstrating substrate utilization during repeated L-36 treatment.

Data are presented as mean ± SEM. Statistical significance was determined as described in the Methods. Unless otherwise indicated, *n* represents independent biological replicates.

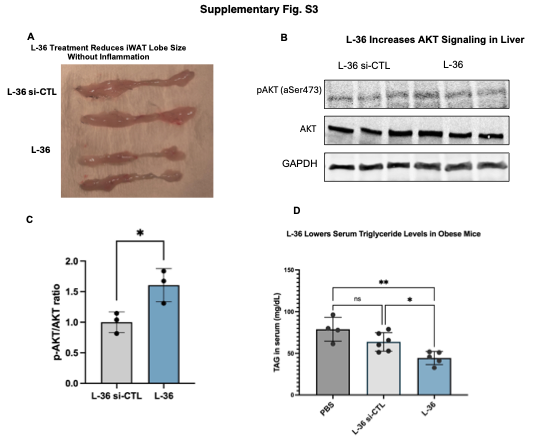

**Supplementary Fig. S3 | Additional histological and biochemical characterization following repeated administration of L-36 LNPs.**

(A) Representative gross images of inguinal white adipose tissue (iWAT) harvested from mice receiving scramble siRNA LNPs (L-36 si-CTL), or L-36 LNPs encapsulating siRNAs targeting TLE3 and ZFP423. (B) Representative immunoblot of phosphorylated AKT (Ser473), total AKT, and GAPDH in iWAT lysates from scramble siRNA-, and L-36-treated mice. (C) Densitometric quantification of the phosphorylated AKT to total AKT ratio (p-AKT/AKT), demonstrating enhanced AKT signaling following L-36 treatment (n = 3). (D) Serum triglyceride (TAG) concentrations measured at the conclusion of the repeated-dosing study (n = 4-6).

Data are presented as mean ± SEM. Statistical significance was determined using unpaired two-tailed Student's *t*-test unless otherwise indicated. Unless otherwise specified, *n* represents independent biological replicates.
