## Supplementary Figure S1 for "Browning Lipid Nanoparticles Deliver Metabolic Benefits via Transient Conversion of White Adipose Tissue"

### Supplementary Fig. S1

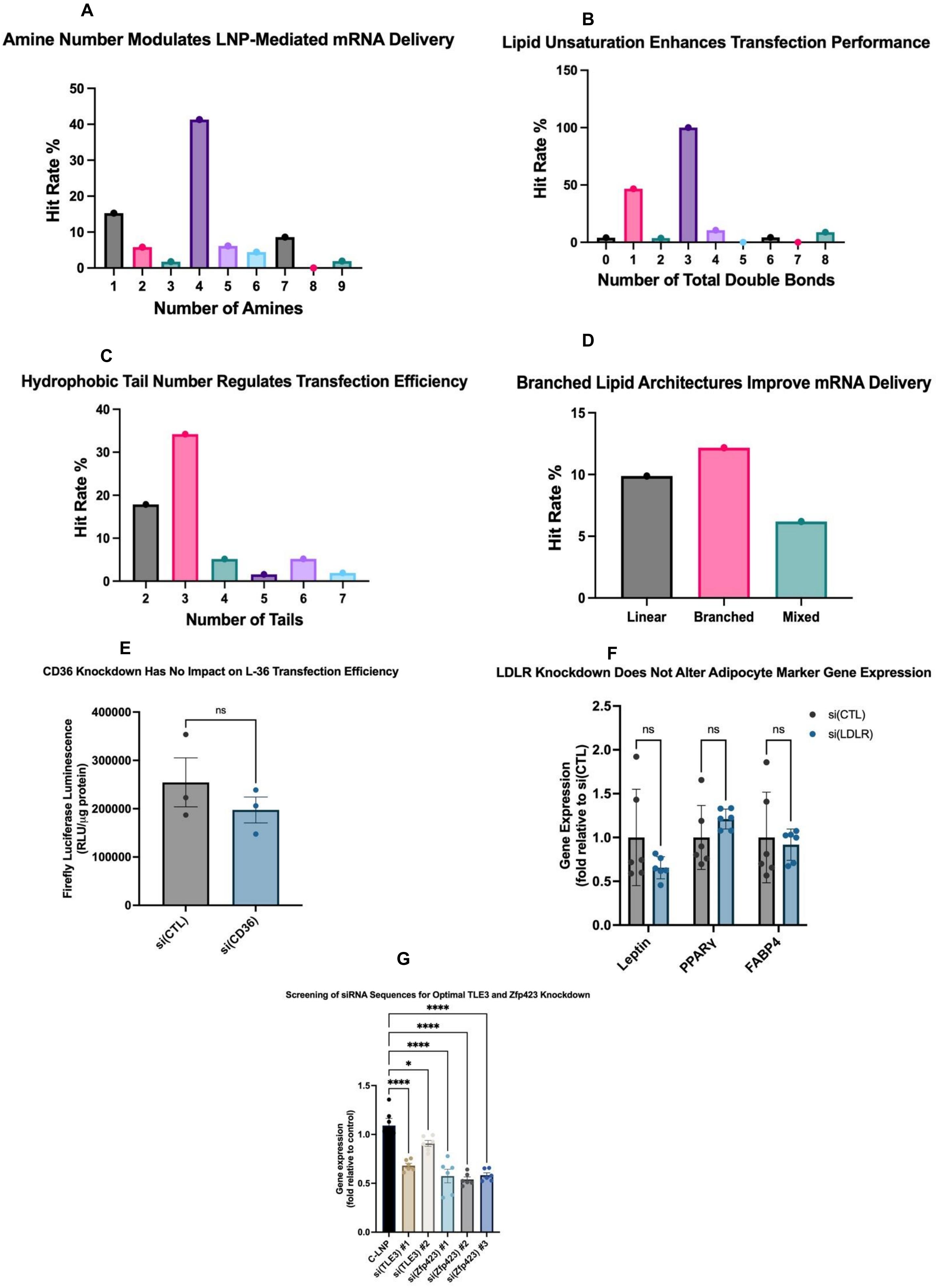

**Supplementary Fig. S1 | Structure-activity relationship analysis identifies key lipid structural features of L-36 and validates experimental controls for mechanistic studies.**
(A) Structure-activity relationship (SAR) analysis showing the effect of the number of amine groups on LNP-mediated mRNA delivery. Lipids containing four amine groups exhibited the highest transfection efficiency, demonstrating that amine number is a critical determinant of LNP performance. (B) Effect of lipid unsaturation on mRNA delivery. LNPs containing lipids with three total double bonds achieved the highest transfection efficiency, indicating that lipid unsaturation strongly influences intracellular delivery. (C) Effect of hydrophobic tail number on transfection efficiency. Lipids containing three hydrophobic tails outperformed alternative architectures, demonstrating the importance of tail number for efficient mRNA delivery. (D) Effect of lipid architecture on transfection efficiency. Branched lipid structures exhibited higher transfection efficiency than linear or mixed architectures, supporting branched lipid design for optimized intracellular delivery. (E) CD36 knockdown in differentiated adipocytes did not alter luciferase expression following L-36 LNP treatment, indicating that CD36 is not required for L-36-mediated cellular uptake or transfection. (F) LDLR knockdown did not significantly alter the expression of adipocyte marker genes, including Leptin, PPARγ, and FABP4, confirming that LDLR silencing does not disrupt adipocyte identity. (G) Screening of Dicer-substrate siRNA sequences targeting TLE3 and Zfp423 in SVF-derived white adipocytes. Transcript levels measured 48 h after treatment identified the most effective siRNA sequences, which were selected for all subsequent in vitro and in vivo studies.
Data are presented as mean ± SEM. Statistical significance was determined using unpaired two-tailed Student's *t*-test or one-way ANOVA with appropriate multiple-comparison correction, as described in the Methods. $P < 0.05$, ****$P < 0.0001$; ns, not significant. Unless otherwise indicated, $n$ represents independent biological replicates.
