## Supplementary Figure S2 for "Browning Lipid Nanoparticles Deliver Metabolic Benefits via Transient Conversion of White Adipose Tissue"

**Supplementary Fig. S2**

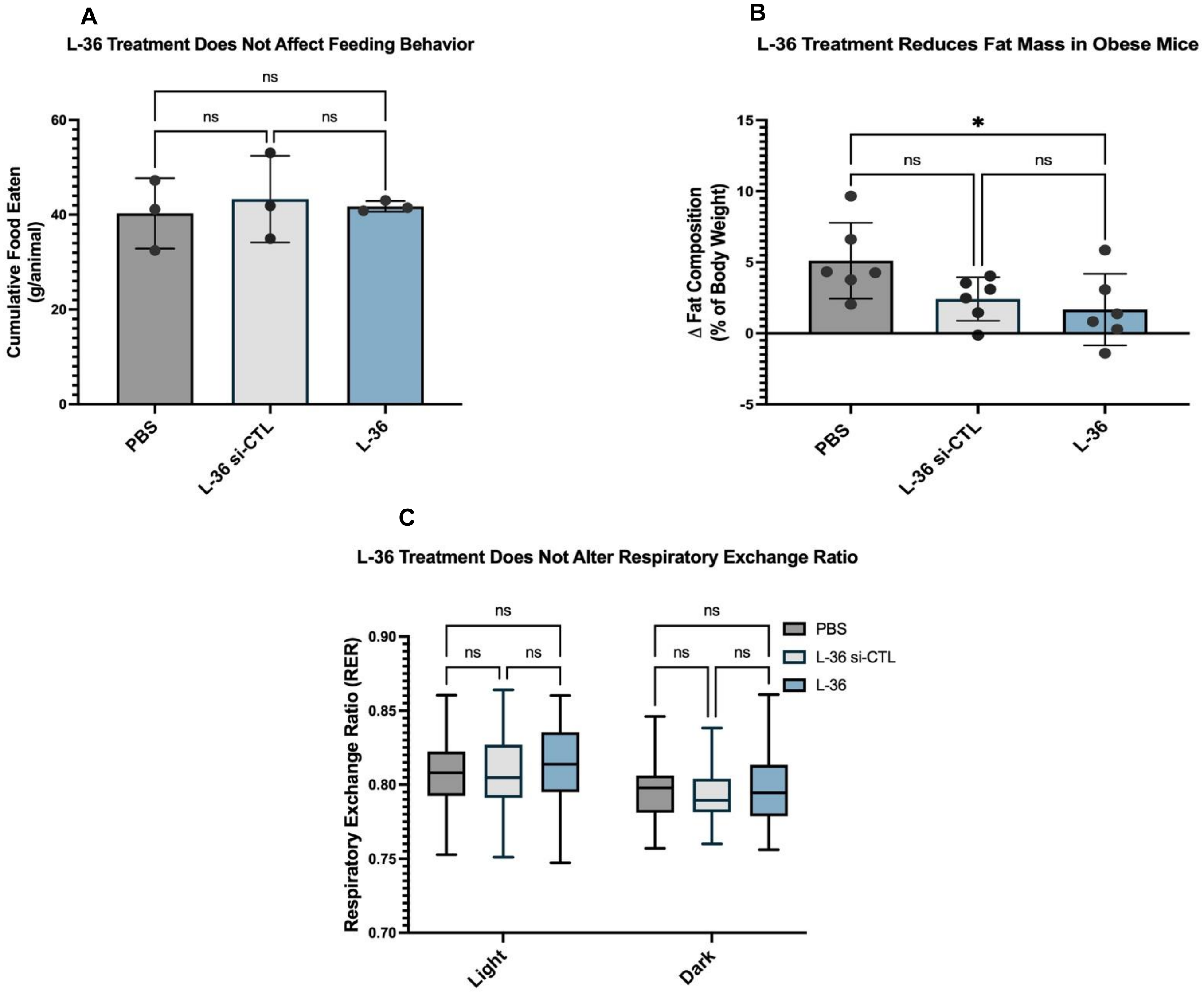


**Supplementary Fig. S2 | Additional metabolic characterization following repeated administration of L-36 lipid nanoparticles targeting TLE3 and ZFP423 in obese mice.**
(A) Cumulative food intake measured throughout the repeated treatment study. Food consumption was comparable between mice receiving PBS or L-36 LNPs encapsulating siRNAs targeting TLE3 and ZFP423, indicating that metabolic improvements were independent of changes in food intake (n = 3 cages per group). (B) Fat mass normalized to pre-treatment baseline following repeated administration of PBS, scramble siRNA LNPs (L-36 si-CTL), or L-36 LNPs encapsulating siRNAs targeting TLE3 and ZFP423. (C) Respiratory exchange ratio (RER) measured by indirect calorimetry, demonstrating substrate utilization during repeated L-36 treatment.
Data are presented as mean ± SEM. Statistical significance was determined as described in the Methods. Unless otherwise indicated, *n* represents independent biological replicates.
