## Supplementary Figure S3 for "Browning Lipid Nanoparticles Deliver Metabolic Benefits via Transient Conversion of White Adipose Tissue"

**Supplementary Fig. S3**

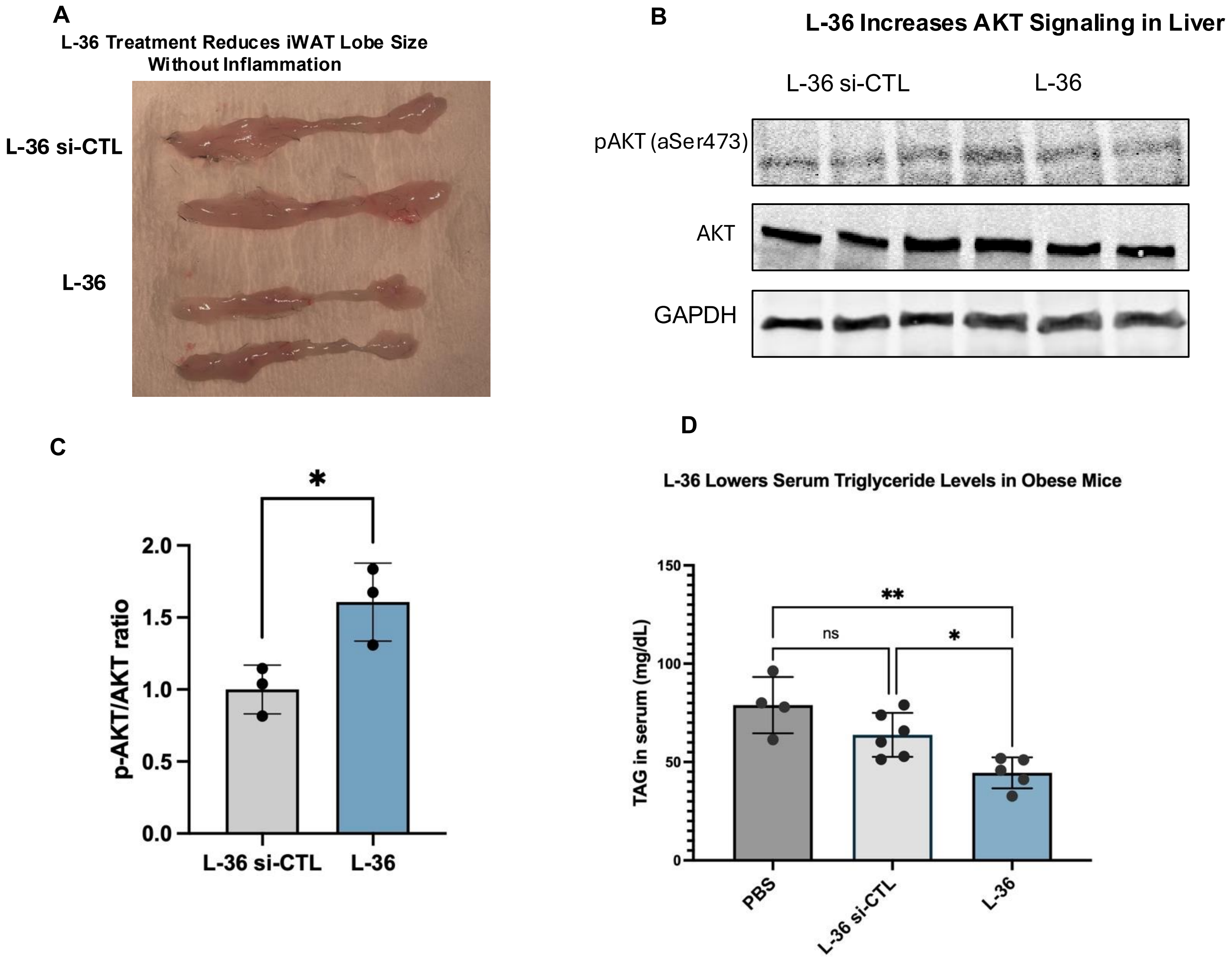


**Supplementary Fig. S3 | Additional histological and biochemical characterization following repeated administration of L-36 LNPs.**

(A) Representative gross images of inguinal white adipose tissue (iWAT) harvested from mice receiving scramble siRNA LNPs (L-36 si-CTL), or L-36 LNPs encapsulating siRNAs targeting TLE3 and ZFP423. (B) Representative immunoblot of phosphorylated AKT (Ser473), total AKT, and GAPDH in iWAT lysates from scramble siRNA-, and L-36-treated mice. (C) Densitometric quantification of the phosphorylated AKT to total AKT ratio (p-AKT/AKT), demonstrating enhanced AKT signaling following L-36 treatment (n = 3). (D) Serum triglyceride (TAG) concentrations measured at the conclusion of the repeated-dosing study (n = 4-6).

Data are presented as mean ± SEM. Statistical significance was determined using unpaired two-tailed Student's *t*-test unless otherwise indicated. Unless otherwise specified, *n* represents independent biological replicates.
